# Maximizing protein production by keeping cells at optimal secretory stress levels using real-time control approaches

**DOI:** 10.1101/2022.11.02.514931

**Authors:** Sebastián Sosa-Carrillo, Henri Galez, Sara Napolitano, François Bertaux, Gregory Batt

**Author notes:** These authors contributed to this work equally (François Bertaux, Gregory Batt).

## Abstract

The production of recombinant proteins is a problem of major industrial and pharmaceutical importance. Secretion of the protein by the host cell considerably simplifies downstream purification processes. However, it is also the limiting production step for many hard-to-secrete proteins. Current solutions involve extensive chassis engineering to favor trafficking and limit protein degradation triggered by excessive secretion-associated stress. Here, we propose instead a regulation-based strategy in which induction is dynamically adjusted based on the current stress level of the cells. Using a small collection of hard-to-secrete proteins and a bioreactor-based platform with automated cytometry measurements, we demonstrate that the regulation sweet spot is indicated by the appearance of a bimodal distribution of internal protein and of secretory stress levels, when a fraction of the cell population accumulates high amounts of proteins, decreases growth, and faces significant stress, that is, experiences a secretion burn-out. In these cells, adaptations capabilities are overwhelmed by a too strong production. With these notions, we define an optimal stress level based on physiological readouts. Then, using real-time control, we demonstrate that a strategy that keeps the stress at optimal levels increases production of a single-chain antibody by 70%.

## Introduction

Bioproduction is a field of major economic importance and is expected to play an important role for the development of a more sustainable industry^1^,^2^. Bio-manufactured products include a variety of chemicals, such as alcohols, organic acids, fragrances, antibiotics, and a large range of industrial or pharmaceutical proteins, such as enzymes and antibodies^3^,^4^. Yeasts are widely used for heterologous protein production. They are inexpensive to grow, easy to engineer, and have extended protein secretory capabilities^5^. This last feature is of great importance, because secreting the proteins of interest (POI) greatly facilitates downstream processes and product purification.

Yet, protein secretion is a complex multi-stage process. A leader peptide acts as a signal for translocation of the protein from the cytosol to the endoplasmic reticulum (ER). There, the protein is folded and undergoes quality control prior to being transported to the Golgi apparatus and being finally secreted^6^,^7^. Bottlenecks may appear at different stages^3^,^8^,^9^,^10^. Secretory stress triggers the activation of various adaptive mechanisms. The unfolded protein response (UPR) plays a pivotal role by modulating the expression of hundreds of genes^11^,^12^,^13^. Its action is twofold. On the one hand, it increases trafficking capacity by regulating genes related to translocation, folding or secretion^11^,^14^. On the other hand, it triggers mechanisms that target the accumulated proteins, such as ER-associated protein degradation (ERAD)^15^,^16^, ER-phagy^17^,^18^ or ER-reflux^19^,^20^. Significant research works have focused on finding genetic modifications of the chassis cells that increase trafficking capacities or that mitigate protein degradation^3^,^8^,^9^,^10^. Unfortunately, such genetic modifications are often chassis- and protein-specific. They are time consuming to identify and implement in yeast cells.

In this study, we propose a fundamentally different strategy. We aim at identifying induction sweet spots, that is, induction levels for which protein secretion is maximal, leveraging adaptation mechanisms that are favorable for secretion and avoiding deleterious ones. We use a small collection of engineered yeast strains expressing various hard-to-secrete POIs under the control of a light-inducible promoter allowing precise control of production demand, and reporting for their UPR secretory stress. We demonstrate that induction sweet spots can be identified by tracking cellular stress levels, and more precisely, their heterogeneity in the cell population. Indeed, a transient bimodality in the POI and UPR levels is observed within the cell population when protein trafficking capacities are overwhelmed and significant protein degradation is triggered. Lastly, we show that secretion levels can be improved by 70% by dynamically keeping the cell population at optimal stress levels using real-time closed loop-control. High sampling frequency and single-cell resolution were essential to obtain these results and were achieved thanks to an open-source automated platform previously developed by our group^21^. The proposed regulation strategy is generic since it applies to any protein to secrete and is complementary with classical chassis-engineering strategies.

## Results

### A systematic experimental strategy for characterizing heterologous protein secretion

Our goal is to study the relations between production demand (ie, induction levels), effective protein production, protein secretion and degradation, and cellular growth for a range of proteins having different secretion complexities. Therefore, we constructed a small collection of yeast strains producing various heterologous proteins. These proteins differ by their posttranslational modifications (PTMs), sizes and native organisms (Supplementary note 1). The approach integrates three main components. The first component relies on a dedicated multi-bioreactor platform with automated cytometry measurements and reactive optogenetic control of yeast in continuous cultures^21^ (Figure 1a). The platform is composed of 8 bioreactors operated in parallel. Each reactor is equipped with a set of LEDs to control gene expression and with an optical density (OD) measurement device. In our experiments, media is renewed to maintain the OD at a target level (turbidostat mode). To ensure a minimal media flow through the reactors, even when the cell growth rate drops due to high secretory burden, we co-cultured an accessory strain having a constant growth rate (Supplementary note 2). Cells in each bioreactor are sampled and measured by cytometry every 45 minutes for several days thanks to a pipetting robot that connects the output line from the reactors to a tabletop flow cytometer. The second component of the approach consists in molecular biology constructs that enable a precise control of the production demand, the measurement of internal levels of the POI together with its secretion-associated stress, and the measurement of secretion levels in the media. The POI is a construct that contains a fusion of the pre-pro-α-factor secretion signal from *S. cerevisiae*^22^,^23^, the protein under study, the mNeonGreen^24^ fluorescent reporter, and 3 copies of the FLAG purification tag (Figure 1b). The fusion of mNeonGreen with the FLAG tags is called the bright tag. Using microscopy imaging, we verified that our bright tag functions as expected, that is, that only the secretion compartments show significant fluorescence in the cell (Supplementary note 3). To control the production of the POI, we use the EL222 optogenetic gene expression system. This light-oxygen-voltage protein is activated by exposure of the cells to blue light^25^. To inform on the stress produced by the secretion of the POI, we took advantage of previous designs using the UPR as a secretory stress reporter by coupling its activation to the expression of a red fluorescent reporter^26^, mScarlet-I^27^. The engineering of our collection of strains is represented in Figure 1c. The third and last component of the approach is a methodology we developed to quantify the secretion levels of the POI in media samples using immunobeads and cytometry (Figure 1d). We use magnetic beads coated with anti-FLAG antibodies to capture the secreted POI (and only the secreted POI). We show that after proper incubation, the fluorescence of the beads is proportional to the quantity of secreted proteins, even at low secretion levels (Supplementary note 3). This experimental setup constitutes a framework to perform the systematic characterization of the secretion process in a generic manner for different POIs (Figure 1e). It allows to conduct parallel experiments over long time scales to obtain dynamic data for different secretion-related processes at single-cell resolution, and for different levels of production demand.

**Fig. 1.**
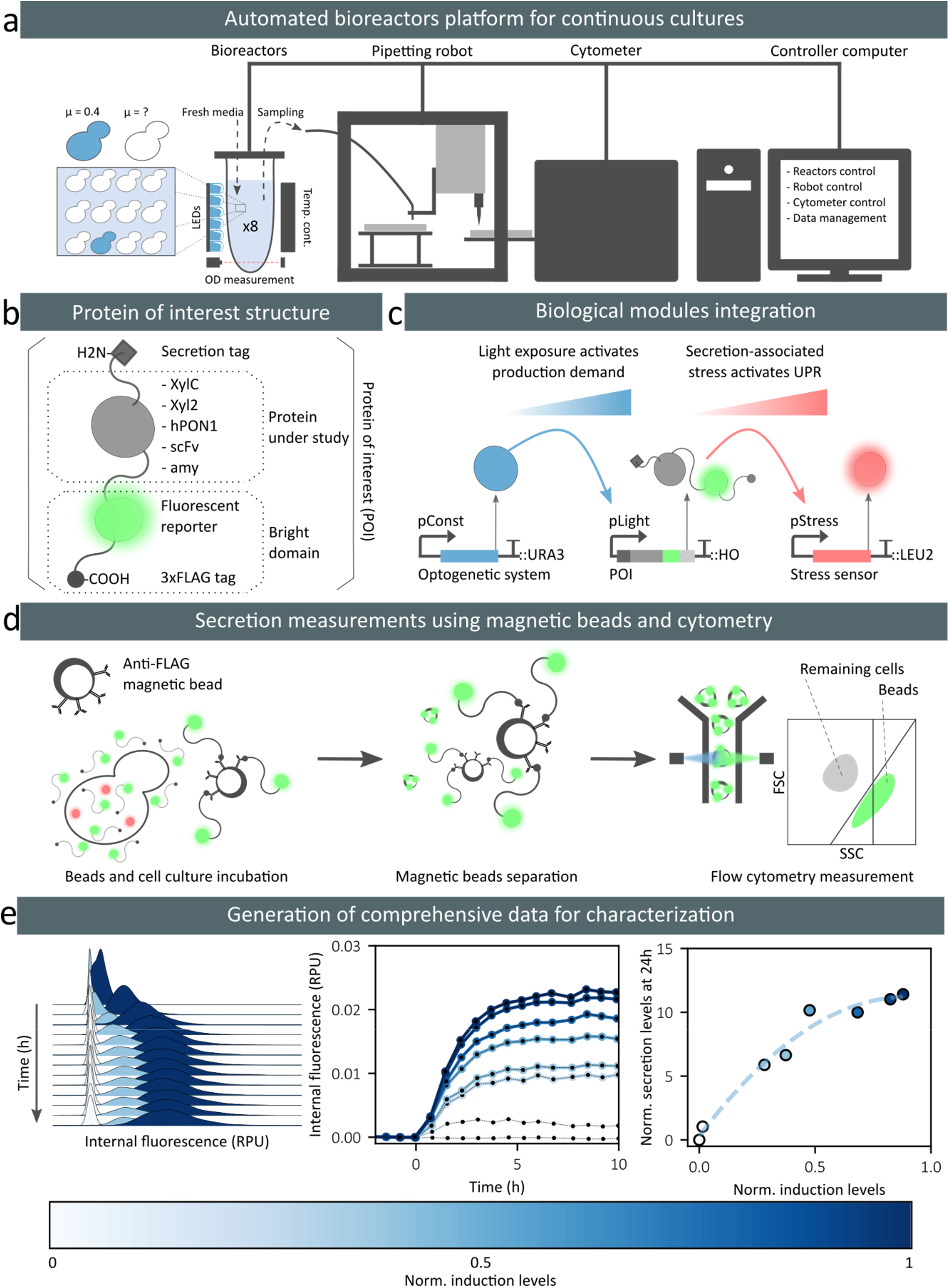
A pipeline for systematic characterization of heterologous protein secretion. **a** The platform uses 8 bioreactors for continuous culture of yeast cells. We co-cultured an accessory strain (blue cells) with the strain of interest (white cells) in a ratio 1:10. The accessory strain is used to assess the growth rate of the strain of interest and the effective light induction levels (Supplementary note 2). It also maintains media influx in the reactor even if the strain of interest is growth arrested (bioreactors are operated in turbidostat mode). The output flow line of the reactors is connected to a pipetting robot that prepares the sample and loads it into the cytometer every 45 minutes. **b** To monitor internal and secreted levels of the heterologous protein under study (Supplementary note 1), the protein is fused to a “bright tag” that comprises of a green fluorescent reporter, mNeonGreen, followed by three copies of the FLAG tag. **c** The biological modules are integrated in the yeast chromosomes. The optogenetic transcription factor EL222 is constitutively expressed (pConst) and activates the pLight promoter upon blue light stimulation. Then, the POI is expressed and may produce secretory-associated stress, activating the UPR response that regulates the expression of genes under the control of the pStress promoter. We introduced a red fluorescent reporter, mScarlet-I, under the control of pStress to monitor the secretory-associated stress. **d** To quantify secretion levels using a cytometer, we developed a methodology based on immuno-magnetic beads capturing only the secreted POI. Beads fluorescence is proportional to the secreted POI levels. The distinction between beads and remaining cells is made thanks to their different light scatter properties (Supplementary note 4) **e** Example of data obtained by the proposed pipeline. RPU: relative promoter units.

### Secretion levels do not correlate with production demand for certain hard-to-secrete proteins

We characterized the impact of heterologous protein secretion on cell physiology and secretory capacity at different production demands and for different hard-to-secrete proteins (Figure 2a). To achieve different levels of production demand using the EL222 optogenetic expression system, we varied the duration of the light exposure within a constant time period of 30 minutes^28^,^29^. Because the light intensities provided by the LED strips might vary from one reactor to another, the effective protein production demand (*ie*, induction level) is evaluated using the co-cultured accessory strain that reports the actual production demand felt by cells. Indeed, it expresses a non-secreted fluorescent reporter under the control of the EL222 transcription factor and integrated into same locus as the POI in the strain of interest.

**Fig. 2.**
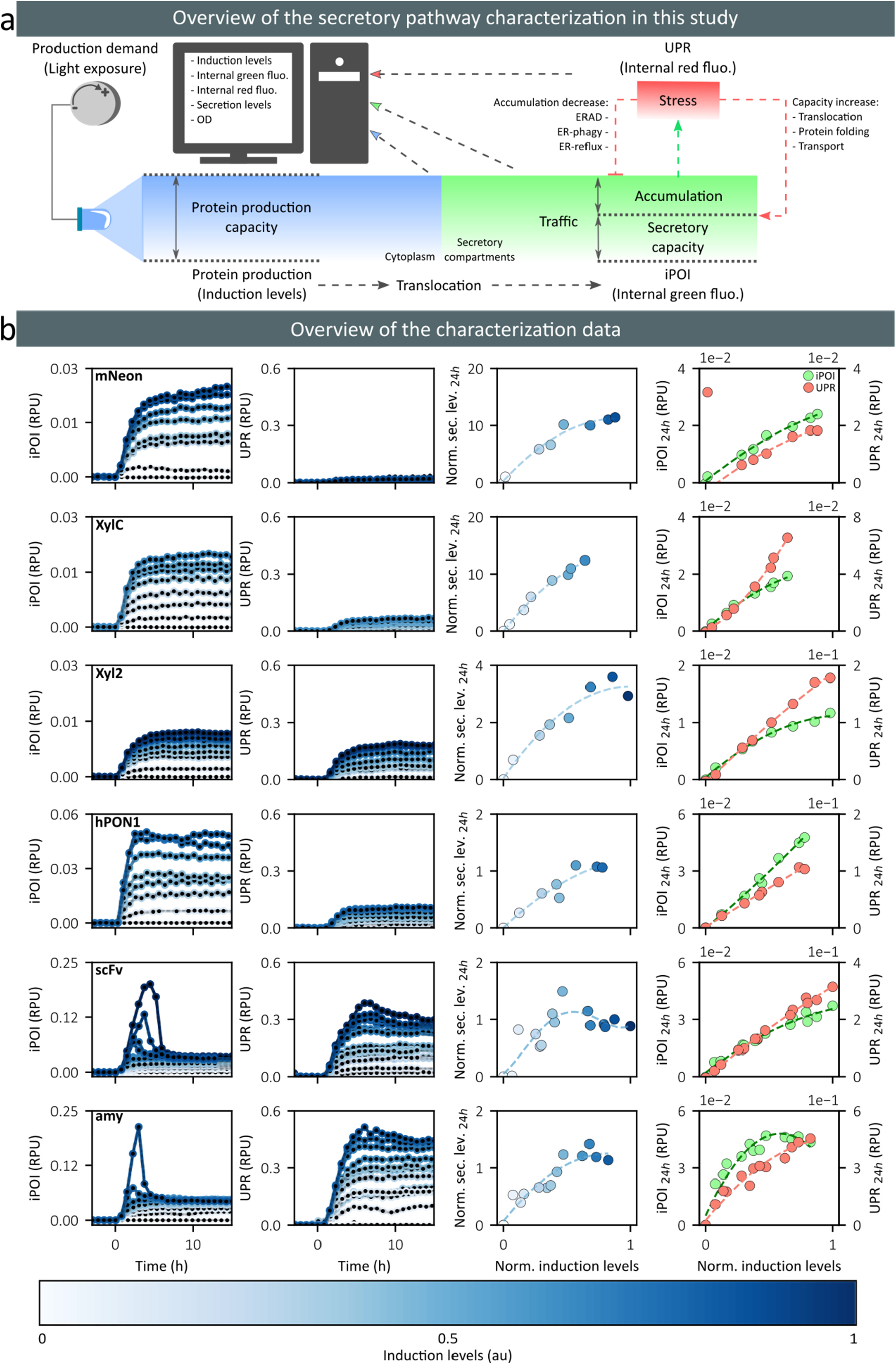
Analysis of protein trafficking and secretion capabilities for a small collection of POIs. **a** Schematic representation of our experimental approach to study protein trafficking and secretion capabilities of cells. The production demand is controlled by blue light illumination of the cell culture. The production of the protein is followed by its translocation into the secretory compartments. Internal protein levels in these compartments (iPOI levels) can be followed by measuring the green fluorescence of cells. The protein can be secreted in the media or can accumulate in the cell. Protein accumulation triggers a secretory stress (UPR stress levels) that can be quantified by measuring the red fluorescence of the cells. The cell stress responses have antagonistic effects with respect to bioproduction: increasing trafficking capabilities or targeting the accumulated proteins for degradation. **b** Characterization of cellular trafficking and secretion capacities for six different POIs. Six strains secreting different POIs were subjected to different production demands. Their iPOI and stress levels were followed in time and secreted protein amounts were quantified at 24h. Rows represent data for the different proteins (indicated in the top left corner of the first column). Columns represent, for different induction strengths, the temporal evolution of the median levels of iPOI distributions, the temporal evolution of the median levels of UPR distributions, the secretion levels at 24h, and the median iPOI (green) and UPR (red) levels at 24h, respectively. Induction levels are color-coded as represented in the bar at the bottom of the figure.

We monitored the iPOI and UPR levels for our six POIs every 45 minutes during 24 hours in continuous culture and under different production demands. Secretion levels were measured at 24 hours. In Figure 2b, we represent the median fluorescence of iPOI and UPR distributions as a function of time. The strain secreting mNeonGreen-3xFLAG (mNeon, Figure 2b, first row) is used to characterize the secretion-associated burden of the bright tag alone. For this strain, we observed that iPOI reaches relatively high levels and then plateaus, whereas the UPR levels remain low. At steady state, iPOI levels and UPR levels increase quasi-linearly with induction strength. Secretion levels are relatively high too. Interestingly, they do not increase linearly with induction strength and iPOI levels, suggesting that secretion rates (and translocation rates) are decreasing at high induction levels. In summary and as expected, mNeon appears to be easy to produce and secrete and is not imposing a significant stress on the cell.

Regarding the Endo-1,4-beta-xylanase C fused to the bright tag (XylC, Figure 2b, second row), the iPOI levels and the secreted POI levels were similar to those observed for mNeon. In contrast the UPR steady state levels were more than doubled. This shows that, in comparison to mNeon, XylC places an higher load on the secretory pathway but that the cell can adapt its trafficking capacities to compensate for this higher load. This is consistent with the fact that XylC requires a disulfide bond for proper folding to pass the pathway quality control (Table S1.1) and that this PTM is catalyzed by the protein disulfide isomerase (Pdi1), whose expression is modulated by the UPR^11^,^30^,^31^.

For the Endo-1,4-beta-xylanase 2 fused to the bright tag (Xyl2, Figure 2b, third row), we observed that the iPOI levels at steady state are half those observed for mNeon and XylC, and that secreted POI levels are comparatively even lower. Moreover, UPR levels are more than twice higher. This indicates that Xyl2 is either poorly translocated or is actively degraded. Relating this behavior with the known PTMs is not obvious here. Indeed since Xyl2 possesses two N-glycosylations, one might have expected that Xyl2 has a longer maturation time than XylC and therefore would accumulate at higher levels.

In the case of the human Paraoxonase 1 fused to the bright tag (hPON1, Figure 2b, fourth row), the data show that iPOI levels at steady state are 5-fold higher than the levels observed for Xyl2, and that secreted levels are half. Given that hPON1 contains a disulfide bond and three N-glycosylations, these observations are consistent with a slow processing of the protein in the secretory compartments in connection with its complex PTM needs.

In contrast to all previously studied proteins, the single chain antibody variable fragment 4M5.3 (scFv, Figure 2b, fifth row) and the α-amylase (amy, Figure 2b, sixth row), fused to bright tags, showed pronounced non-monotonic dynamics for their iPOI levels as well as for their UPR levels. Peaks in iPOI and stress levels are observed a few hours after strong inductions. Moreover, scFv and α-amylase-secreting strains showed the highest stress levels (10 to 15 times more than mNeon at steady state). The burden generated by the secretion of these POIs is also evident when looking at the growth rate dynamics. After three hours of induction the growth rate decreases up to 40% of the pre-induction levels and then recovers (Supplementary note 5). Importantly, for the strain secreting scFv, the relation between induction and secretion levels is non-monotonic. This shows the presence of an optimal level of external demand that maximizes protein secretion. From the perspective of protein bioproduction, this secretion sweet spot is important since further increasing the production demand is counter-productive for protein production. The situation appears to be slightly better for the strain secreting the α-amylase (Figure 2b, sixth row). We observed a saturation of the secretion levels for induction above 50% of the maximum but no marked decrease. Yet stress levels continue to increase, showing that high induction levels cause unnecessary stress on the cells.

In summary, this dataset shows very different secretion behaviors for the different POIs, confirming that secretory constraints and bottlenecks are indeed protein-specific. In particular, we observed induction sweet spots above which cellular secretory stress increases and protein production plateaus or even decreases. Such situations are associated with surprising non-monotonic dynamics in iPOI and stress levels.

### Single-cell data reveal a state of secretion burnout in a fraction of the cell population

A closer analysis to cytometry data reveals that tracking median levels only can be misleading. Indeed, the peak of internal protein levels for scFv and amylase-secreting cells corresponds to a bimodal population. A fraction of these cell populations transiently accumulated iPOI at much higher levels than the other cells (Figure 3a). From now on, these cells will be referred to as “accumulators”. Formally, iPOI-accumulators, or more simply accumulators are defined as having iPOI levels higher than the mean plus three standard deviations of a normal distribution fitted to the cell population at steady state. The presence of the accumulator cells is correlated over time with a decrease in the overall growth rate, strongly suggesting that these cells have a reduced growth rate (Supplementary note 5). This observation is consistent with previous studies describing that in ER-stressed cells, a minimum level of ER functionality is required to complete cell division. When cells recover by the action different adaptive responses, ER capacity and growth rate are restored^32^,^33^. After a delay, the distribution of UPR levels also appears to be bimodal. When studying the abundance of accumulator cells, defined as the highest fraction of accumulators over the course of the experiment, we saw that their abundance increases proportionally to the induction levels above a given threshold. Moreover, the level of iPOI in these accumulator cells is also increasing linearly with the induction level, above a given threshold (Supplementary note 6). This strongly indicates that the secretory pathway and the degradation pathway are both saturated in accumulator cells. In summary, when proteins are produced at high levels and cells do not adapt rapidly enough, proteins accumulate in the secretory pathway, overwhelming cell adaptation capacities and impacting growth. Growth-arrested cells are then likely to accumulate proteins even further. These cells are experiencing a secretion burnout. With time, they might adapt and recover growth.

**Fig.3.**
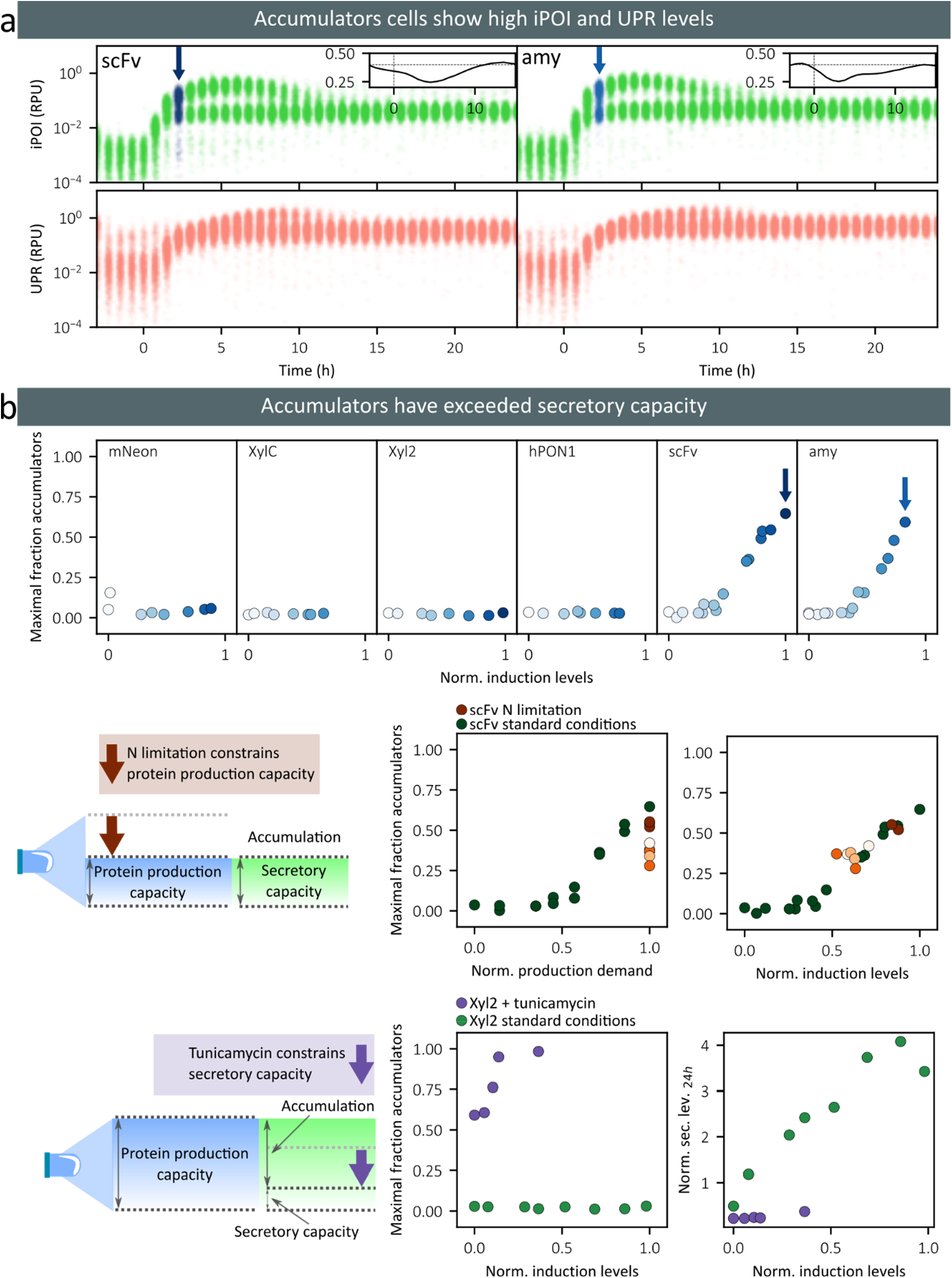
Accumulators signal that secretory capacities are overflown. **a** Single-cell data for iPOI (first row, green dots) and UPR (second row, red dots) at maximal induction levels for scFv (first column) and amylase (second column). Blue arrows indicate when the fraction of accumulators is maximal (see Figure 3b too). Insets represents the temporal evolution of cell growth rate when accumulators are present. The horizontal dotted line indicates the pre-induction growth rate (0.4 h^-1^). **b** At the top, we show the maximal fraction of accumulator cells observed over the course of experiments as a function of induction levels. Maxima always appear between 2 and 4 hours after induction. The intensity of the blue is proportional to the induction levels. In the center, we show the data obtained in the experiment in which the protein production capacity was constrained by limiting the nitrogen supplied using the scFv-secreting strain. The maximal fractions of accumulator cells quantified in nitrogen-limited conditions and in normal conditions are shown in orange and green, respectively. The intensity of the orange color corresponds to the ammonium sulfate concentration supplied in each conditions (0, 5, 50, 500 or 5000 mg/ml). The left and right plots represent the maximal fraction of accumulator cells as a function of the production demand and as a function of the induction levels, respectively. At the bottom, we show the data obtained in the experiment in which the secretory capacity was limited by adding tunicamycin in the media of the Xyl2-secreting strain. The maximal fractions of accumulator cells in tunicamycin and in normal conditions are shown in purple and green, respectively. The left and right plots represent the maximal fractions of accumulator cells and the secretion levels measured at 24 hours as a function of induction levels, respectively.

The appearance of accumulators cells is therefore a sign that maximal trafficking capacities have been reached. To test the genericity of this observation, we grew cells in different contexts. Firstly, we constrained nitrogen availability in the culture medium, thereby limiting cellular capacities to produce proteins^34^,^35^. For scFv-secreting cells, we observed that nitrogen limitations strongly reduce the proportion of accumulators in the cell population (Figure 3b, middle). Moreover, thanks to the accessory strain, that is also growing in nitrogen-limited conditions, we can estimate the effective protein production rate in these conditions. We found that in normal media or in nitrogen-depleted media, the same protein production rate leads to the same fraction of accumulator cells. Secondly, we added tunicamycin to the media, a drug that inhibits the N-glycosylations catalyzed in the ER, thereby decreasing the trafficking capacities of the secretory pathway in cells. For Xyl2-secreting cells, we observed that in presence of tunicamycin, accumulators are present even in absence of induction and that their number rapidly increases with induction levels (Figure 3b, bottom). In summary, we observed for different proteins (ScFv, Amylase and Xyl2) and in different conditions (normal media, low nitrogen and presence of tunicamycin) that the presence and the relative fraction of accumulators signals the existence and the severity of trafficking issues in the secretory pathway.

### UPR-mediated adaptation is essential to prevent and recover from secretion burnout

When confronted to a too strong production demand, cells accumulate proteins to extremely high levels, stop growing, and experience high stress levels. Yet, this accumulator phenotype is transient and after approximatively 5 hours, protein levels and stress levels decrease and with time reach the levels of the population of non-accumulator cells. Here, we investigate the role of the stress adaptation pathway in this rather spectacular adaptation. To do so, we constructed Hac1 deficient strains. Hac1 is the transcription factor that binds to the promoter controlling the expression of UPR-regulated genes^11^,^36^. Therefore, Hac1-deficient cells are not capable to trigger the UPR.

The *HAC1* knockout mutant of the scFv-secreting strain showed a very severe phenotype. The entire cell population shows the accumulator phenotype, stop growing, does not recover and gets eventually diluted away from the bioreactor (Figure 4). This observation demonstrates that UPR-mediated adaptation is the principal, if not unique, mechanism of adaptation for cells experiencing secretion burnout. More surprisingly, accumulators also appeared for the *HAC1* mutant of the mNeon-secreting strain. Their fraction is low at low induction levels but rapidly increases with increasing induction levels. This reveals that in *S. cerevisiae*, the nominal trafficking capabilities are exactly matching the needs of the native secretome and that any additional demand necessitates some adaptation.

**Fig.4.**
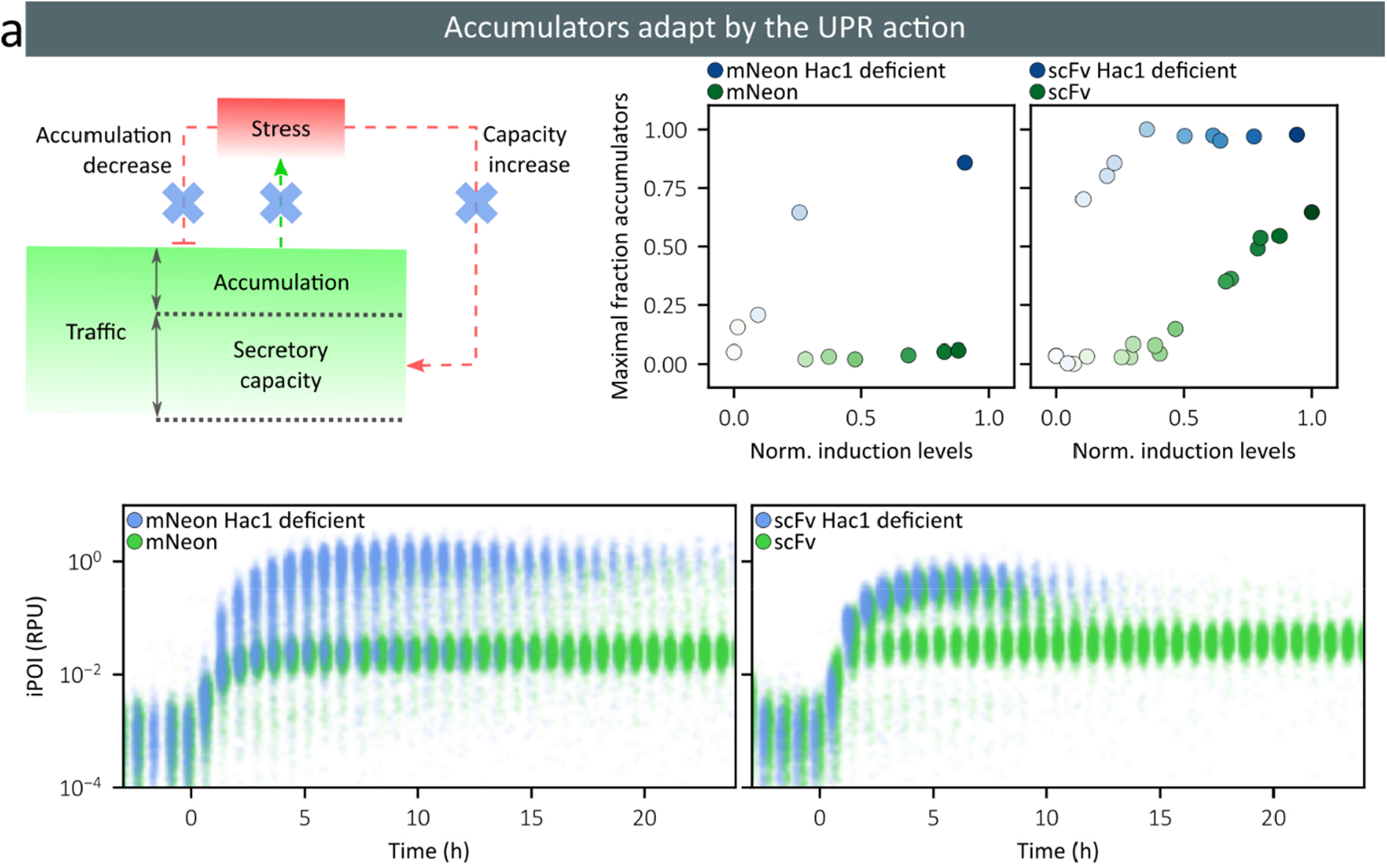
UPR-mediated response is essential for the adaptation of accumulator cells and for maintaining physiology at all induction levels. At the top, we show the maximal fractions of accumulator cells for Hac1 deficient cells (blue) and for normal cells (green), as a function of the induction levels (left: mNeon-secreting cells, right: scFv-secreting cells). The intensity of the color is proportional to the induction levels. At the bottom, we show single-cell data for four representative characterization experiments with maximal induction levels (blue dots: knockout strain, green dots: wild-type strain; left: mNeon-secreting cells, right: scFv-secreting cells).

### The appearance of accumulators signals the activation of the degradation pathway in the entire cell population

We have previously identified that some hard-to-secrete proteins, namely scFv and amylase, have induction sweet spots (Figure 2). Their efficient production necessitates that the induction of gene expression is made at carefully chosen levels. However, scanning induction levels and quantifying secreted protein levels in a systematic manner is time consuming. In Figure 5a, we show that secretion sweet spots can instead be identified by tracking the appearance of accumulators in the cell population. Indeed, the induction levels that maximize protein production correspond to the appearance of accumulators, where appearance is defined as an increase of more than 5% above basal level. Note that we define here accumulators with respect to UPR levels instead of iPOI levels (bimodal distributions are observed in both cases). This definition is more convenient here since this criterion is protein independent and does not require complex genetic constructions. The maximal fractions of iPOI-defined accumulators and of UPR-defined accumulators are linearly related (Supplementary note 6).

**Fig. 5.**
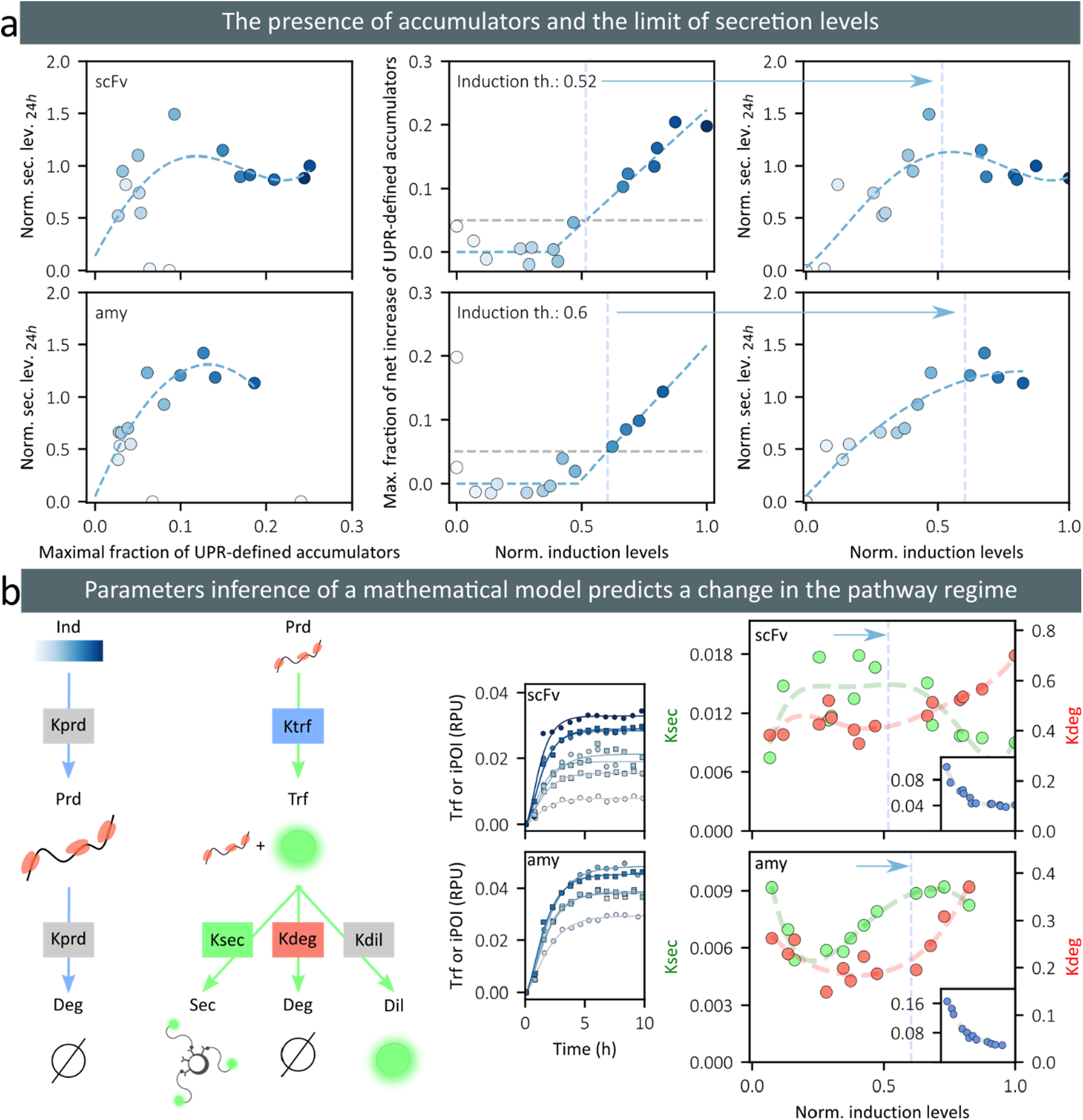
The appearance of accumulator cells indicates secretion sweet spots where the balance between protein secretion and degradation is optimal. **a** On the left, the plots show the relationship between the occurrence of (UPR-)accumulator cells and the protein secretion levels at 24h (top: scFv-secreting strain, bottom: amylase-secreting strain). In the middle, the plots represent the maximal fraction of UPR-accumulators as a function of the induction levels. The induction thresholds indicate the levels of induction at which the net increase of UPR-accumulator cells reaches 5% of the population (blue dotted bars). On the right, the plots represent the secretion levels at 24h as a function of the induction levels (same as in Figure 2b), with an indication of the previously defined induction thresholds (blue dotted bars). We observe that they correspond to maximal secretion levels. The intensity of the blue color corresponds to the induction levels. **b** On the left, we provide a schematic representation of our simple mathematical model showing the production and translocation of the protein in the trafficking compartments and its possible outcomes: secretion, active degradation, or dilution due to cell growth. Parameters in grey boxes are calibrated using independent experiments and have fixed values. The other parameters are calibrated using time series and secretion data, independently for each induction level. In the middle, the plots represent the temporal evolution of the iPOI levels in the non-accumulator cell population together with model fits (top: scFv-secreting strain, bottom: amylase-secreting strain). On the right, the plots represent fitted parameter values as a function of induction levels (Ksec: green, Kdeg: red, and Ktrf: blue in inset; top: scFv-secreting strain, bottom: amylase-secreting strain). Blue dotted bars indicate the previously defined induction thresholds.

We developed a simple mathematical model to better understand the cell physiology around these production sweet spots. We focused on better understanding the relative impact of the adaptative responses that increase trafficking capacities or that target proteins for degradation. Distinguishing these antagonistic effects is essential for understanding production efficiency. Because the accumulator population is marginal around secretion sweet spots and accumulators are present only transiently, we focused on understanding the adaptation response of the non-accumulator cells. The model is based on ordinary differential equations and captures three main processes with simple assumptions. A protein production intermediate, Prd, is produced proportionally to the optogenetic induction level and is degraded at a rate proportional to its concentration. This intermediate can typically be the messenger RNA of the POI. It is important to account for the initial delay of protein appearance. This Prd intermediate is used to produce a protein that is translocated in the secretory compartments. This protein in traffic, Trf, is then either secreted, actively degraded (by ERAD for example), or diluted in the cell because of cellular growth. The model also accounts for the secretion of the proteins by cells in the media, and their dilution due to media renewal at a rate that equals the cell growth rate (turbidostat mode). To simplify parameter identification, the model is first fitted to experimental data obtained for a strain expressing a non-secreted mNeonGreen protein (Ksec=Kdeg=0). We obtained parameters values for Kprd and Kdil that were consistent with our experimental conditions and with literature values^37^,^38^ (mRNA half-life of 20 minutes and cell generation time of 90 minutes). The model was then independently fitted to each time-series to identify how the parameter values evolve as a function of the production demand. The model, together with the data for the non-accumulator cell population, model fits, and the corresponding parameter values are represented in Figure 5b. For both POIs, the parameter estimates show that protein degradation rates increase when induction exceeds the accumulator-appearance thresholds defined previously. At high induction levels, protein secretion rates either plateau (amylase) or decrease (scFv). In summary, our quantitative analysis reveals that at the induction levels where accumulators appear, cells have reached their maximal trafficking adaptation capabilities. Increasing further the demand leads to increasing protein degradation in the non-accumulator cells or transiently experiencing a secretion burnout in accumulator cells.

### Maximizing protein production using real-time control and optimal stress levels

The previous findings can be used to propose a real-time control strategy to maximize protein bioproduction. Firstly, one identifies the optimal level of stress, that is the UPR stress level that is associated with the apparition of accumulators. This should indicate a physiological state in which cell adaptation is optimal for secretion. As before, we define this target stress level as the stress felt by cells when light induction was such that the maximal fraction of accumulators is 5% above baseline levels (Figure 6a). We use mean stress levels instead of median values so that our control strategy can be implemented using simpler measurement devices than cytometers, such as plate-readers for example. We then propose a very simple control strategy. When UPR levels were below the reference level, the duration of light stimulation is increased by a fixed amount (5% of sampling period), and conversely, when UPR levels were above the reference, the duration of the light stimulation is reduced by the same amount. We start with no light. Here, we take advantage of the real-time control capabilities of our experimental platform, which has been previously employed to achieve real time control in diverse experimental contexts^21^,^39^,^40^. Several real-time control experiments were performed using reference UPR levels in the vicinity of the previously-defined target level, and protein production by scFv-secreting cells was quantified in the media. We found that secretion levels were indeed optimal when the effective UPR levels were maintained close to the target level (Figure 6b). Moreover, we found that this optimal level is 70% higher that the level of protein one obtains using a constant, full-light induction (Figure 6b). This experiment demonstrates that by tracking the apparition of accumulator cells, one can define a target stress level that leads to optimal protein production.

**Fig. 6.**
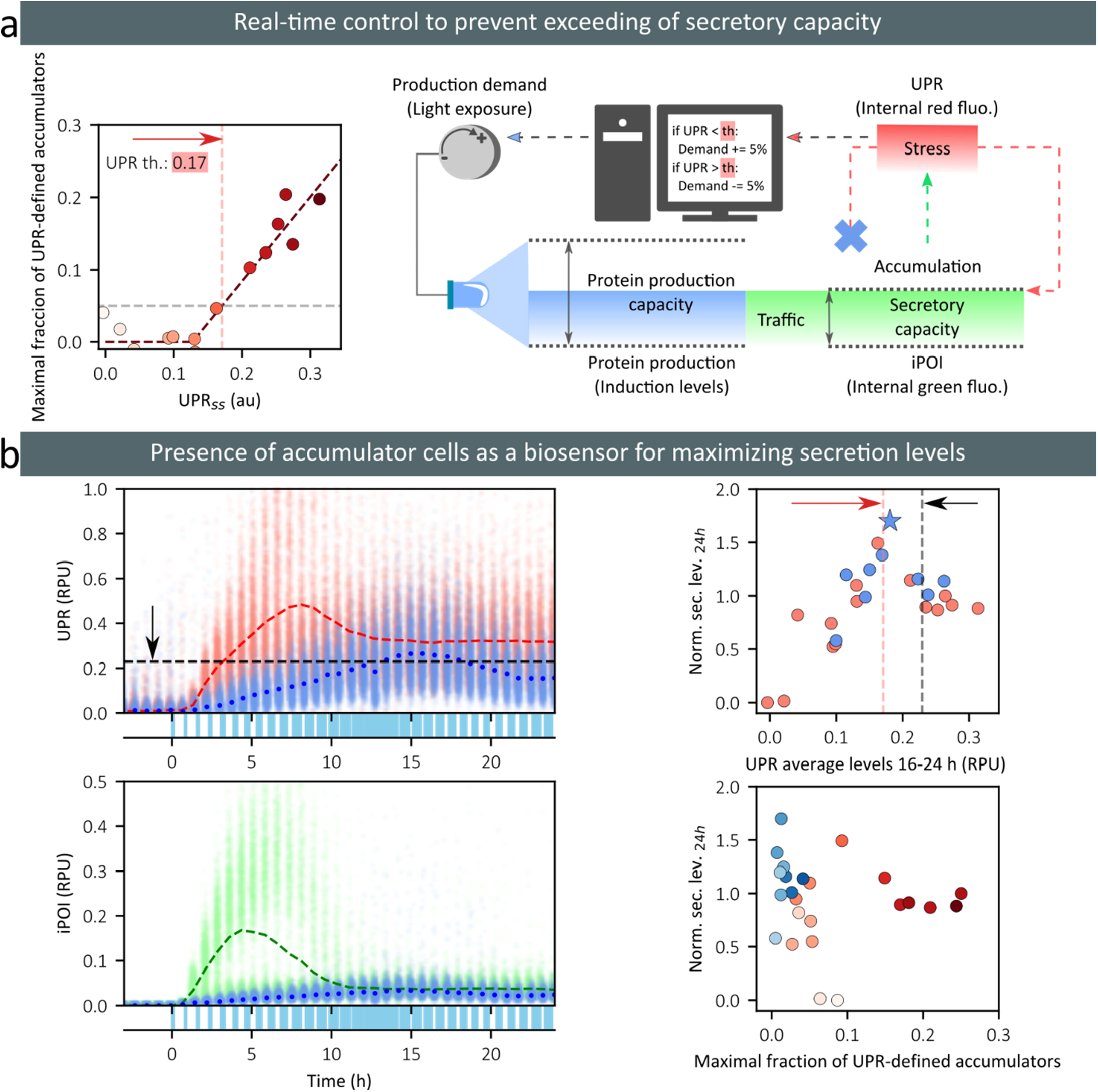
Real-time control approaches that keep cells at optimal stress levels maximize secreted protein levels. **a** On the left, the plot shows the maximal fraction of UPR-defined accumulator cells as a function of the mean stress level of the population at 24h (the data comes from the characterization experiments represented in Figure 2b). The target stress level is defined as the UPR level at which the net increase of UPR-accumulator cells reaches 5% of the population. On the right, we provide a schematic representation of our control strategy that aims at maximizing the productive adaptations and minimizing the deleterious adaptations of the cell to secretory stress. The average levels of UPR are monitored every 45 minutes, and depending on their values with respect to a reference value (th), the production demand is increased or decreased by 5% of the stimulation period (45 minutes). **b** On the left, the single-cell data for UPR (top) or iPOI (bottom) levels are represented as a function of time in a full-induction experiment (red and green dots) or in a real-time control experiment (th=0.23, light blue dots). Mean values are represented by dashed (red and green) lines for the full-induction experiment, and by darker blue dots for the real-time control experiment. The light stimulus for the real-time control experiment is represented at the bottom of the plots (blue rectangles). In this experiment, the average of the UPR levels during the last 8 hours of experiment equals 0.17. On the right, we represent the secretion levels obtained in different real-time control experiments (blue dots) and in different characterization experiments (red dots) as a function of the average UPR levels over the last 8 hours of the experiment (top) or as a function of the maximal fraction of UPR-defined accumulators (bottom). The blue star corresponds to the secretion levels of the real-time control experiment shown on the left. In the top plot, the target level (ie the optimal stress level defined in the panel above) and the reference level for the experiment shown on the left are shown by red and black bars, respectively. In the bottom plot, the intensity of the color corresponds to the average UPR levels over the last 8 hours of experiment.

## Discussion

We presented a systematic approach to characterize yeast trafficking and secretory capacities for a panel of proteins with different structural features. We generated dynamic data at the single-cell resolution over a range of production demands. Their analysis allowed us to discover that a subpopulation of cells accumulates the POI, and that the fraction of accumulator cells increases with induction levels. We showed that these cells have exceeded their maximal trafficking capacities. These cells accumulate proteins to very high levels, stop growing which worsen the situation and leads to high stress levels. They experience a secretion burnout. We also show that in these situations, cell adaptation is largely, if not exclusively, mediated via the UPR response. Importantly, the presence of accumulator cells is a single-cell phenomenon that reflects a population effect. Indeed, the emergence of accumulator cells signals that maximal trafficking capacities are reached by the cell population and that increasing further the demand only leads to creating more accumulator cells or to activating more the protein degradation adaptative response in non-accumulator cells. This explains why the emergence of accumulators signal a protein production sweet spot. This is strongly supported by the fact that protein production was found to be maximal in real-time control experiments that aim at tuning the demand to maintain the cellular stress at the level at which accumulators appear. Notably, secreted levels were found to be 70% higher than when the demand was maximal. We note that the presence of bimodal distributions for secretion-associated features in yeast has already been described in previous works using microfluidics platforms^41^. Yet, their significance for optimizing protein production was not discussed. Similarly, other studies have already reported evidence for the existence of an optimal level of demand to optimize secretion^42^,^43^. Yet, no criteria were then provided to identify such production sweet spots using sensors of the cell physiology.

The secretory pathway is a very complex cellular process^6^,^11^. The optimization of protein production has proven to be extremely challenging^3^,^9^,^42^. Yet our approach can be applied in principle to improve the secretion of any protein in any yeast or more generally eukaryotic organism since the UPR sensor is a very generic reporter. This is of major relevance because each protein generates specific bottlenecks that may also depend on the host strain, optimized or not. Therefore, we provide a generic approach for optimizing protein production that is complementary to other strain engineering approaches.

## Methods

### Cloning and strains construction

The genetic constructions used in this study were designed to be compatible with modular cloning strategy *Yeast Tool Kit* developed by Lee and colleagues^44^. Some parts corresponding to coding sequences or promoters were purchased from external suppliers, Twist Biosciences or GENEWIZ, depending on the specific needs. The coding sequences of the proteins of interest were obtained from literature and verified in *GeneBank*^45^ (Supplementary note 1). The sequences were codon optimized for *S. cerevisiae* using IDT (Integrated DNA Technologies) codon optimization tool. In all cases the final vectors are integrative plasmids for yeast (Supplementary note 1). Transformation were based on an adapted version of the lithium acetate/single-stranded carrier DNA/PEG method of transformation from Gietz and colleagues^46^,^47^. All strains are derived from the common laboratory strain BY4741. Strain details are described in Supplementary note 1. All strains used in this work express the light-inducible transcription factor EL222 from the *URA3* locus (transcriptional unit: pTDH3 NLS-VP16-EL222 tSSA1). In all cases, the POI expression cassettes are integrated in the *HO* locus, and confers constitutive expression of *HIS3* for positive selection. The transcriptional unit includes the light inducible promoter, composed of five copies of the pC120 EL222-binding sequence and the minimal promoter CYC180, followed by the POI coding sequence (transcriptional unit: pC120×5-pCYC180 POI tTDH1). The ER-associated stress reporter is adapted from Pincus and colleagues^26^, integrated in the *LEU2* locus (transcriptional unit: UPREx4-pCYC180 mScarlet-I tENO1). To generate the *HAC1* knockout, we used a CRISPR/Cas9-based^48^ method to introduce an early 5’ TAG amber STOP codon replacing the protospacer adjacent motif (PAM) within the coding sequence of the target gene. All strains used in this study were sequenced for the proper integration of expression cassettes.

### Culture conditions for characterization experiments

All characterization experiments were performed in 20 mL of culture volume in the bioreactors at 30°C, in turbidostat mode (OD 0.5, typically corresponding to 10^7^ cells/mL according to cytometry data). The media used was synthetic complete (Formedium LoFlo yeast nitrogen base CYN6510 and Formedium complete supplement mixture DCS0019), 2% glucose (w/v) and 5 mM of L-arginine (Sigma 11009). L-arginine is used to maintain pH at 7, thus keeping secreted protein integrity along the experiment duration to allow secretion measurements^49^,^50^,^51^. After an overnight (16 hours approx.) within the bioreactors, the automated cytometry measurements started. Samples were taken every 45 minutes and 5000 events were recorded by cytometry. Approximately after 3-4 hours of measurements in dark conditions, the LEDs of the different reactors were switched on for various durations within periods of 30 minutes during 24 hours (LED intensity was 20). To provide reliable and accurate measurements of the growth rate and induction levels, as well as to keep media flow through the reactors, a yeast accessory strain was co-cultured with the strain of interest at an initial ratio of 1:10 in all experiments (Supplementary note 2). All experiments were protected from direct light.

### Culture conditions for nitrogen-limiting experiment

For the nitrogen-limiting experiment the media used was synthetic complete without ammonium sulfate (Formedium LoFlo yeast nitrogen base without ammonium sulfate CYN6210 + Formedium complete supplement mixture DCS0019), 2% glucose (w/v) and 5 mM of L-arginine (Sigma 11009). Then, the ammonium sulfate was added (Sigma ammonium sulfate A4418) to reach final concentrations of 5, 50, 500 and 5000 mg/ml. Cytometry sampling frequency and settings, LEDs intensity, use of the accessory strain, temperature, and culture volume were similar to the characterization experiments. In this experiment, the OD was dynamically maintained between 0.4 and 0.6. The culture was kept growing during an overnight in synthetic complete media, 2% glucose (w/v) and 5 mM of L-arginine. Then, we change to nitrogen-limiting media and started cytometry measurements. Eight hours after, the optogenetic induction with constant light was triggered during 24 hours.

### Culture conditions for tunicamycin stress-induced experiment

In the tunicamycin stress-induced experiment all settings were similar to those used in the characterization experiments. The media used was synthetic complete, 2% glucose (w/v) and 5 mM of L-arginine, with the addition of tunicamycin in DMSO (Bio-Techne 3516) at 0.25 mg/ml final concentration. The culture was kept growing during an overnight in synthetic complete media, 2% glucose (w/v) and 5 mM of L-arginine. Then, we change to tunicamycin containing media and started the cytometry measurements. Eight hours after, the optogenetic induction with constant light was triggered during 24 hours. Previous characterization experiments were done to decide on the concentration of tunicamycin and the DMSO effects on the cell culture.

### Culture conditions for control experiments

In the control experiments all settings and media were similar to those used in the characterization experiments. The accessory strain was not used (with one exception) because we wanted to minimize the probability of counting red fluorescence from the sensor strain as UPR signal. There were three sets of experiments produced in different days, and each of them included one in which maximal demand strategy was applied. Such experiment was used as reference for secretion levels normalization and quantification of maximal fraction of accumulators in the given experiment set. In all experiment sets, after an overnight of growth within the bioreactors, the automated cytometry measurements started sampling every 45 minutes. Approximately after 3-4 hours of measurements in dark conditions, the LEDs of the platform were switched, at intensity 20, with duty cycles set by the control strategy. Note that in feedback experiments, the light stimulation period was of 45 minutes to match the sampling period.

### Accessory strain for assessing effective induction levels and quantifying growth rates

To provide reliable and accurate measurements of the growth rate and induction levels, a yeast accessory strain was co-cultured with the strain of interest at an initial ratio of 1:10 (Supplementary note 2). To allow differentiating one yeast strain from the other in co-cultures, the accessory strain constitutively expresses a blue fluorescent reporter (transcriptional unit: 2x[pTDH3 mCerulean tTDH1], integrated in the *LEU2* locus and conferring *LEU2* constitutive expression). Moreover, the accessory strain expresses a cytoplasmic red fluorescent protein, whose expression is controlled by the EL222 optogenetic system as for the gene of interest in the strain under study (transcriptional unit: pC120×5-pCYC180 mScarlet-I tTDH1, integrated in the *HO* locus and conferring *HIS3* constitutive expression). Therefore, at steady state, expression levels of the red fluorescent protein in the accessory strain inform on the production levels of the POI in the strain of interest. In our data, the effective induction levels are inferred by normalizing the red fluorescence of the accessory strain to its maximal value across all experiments shown in this study. Moreover, we use the accessory strain to assess growth rate of the strain of interest. Indeed the dynamics of the relative amount of the two strains allows us to infer the growth rate difference for each condition (Supplementary note 2).

### Cytometry data analysis

To account for day-to-day variability and settings adjustments for the cytometer (Guava EasyCyte 14HT, 362 Luminex), we computed a correction coefficient for all the channels used in each experiment. Such coefficient was computed by using the *Guava*^®^ *easyCheck™* Kit, in which fluorescent beads are measured in every channel, providing the changes in mean fluorescence intensities from one experiment to another. These coefficients are then used to normalize the fluorescence obtained from the different channels in each experiment. The channels used for each fluorescent reporter were mNeonGreen: GRN-B channel, mCerulean: BLU-V channel and mScarlet-I: ORG-G channel. Gating is used on the forward light scatter (FSC) channel to discard cell doublets and cell debris (only events having values between 1000 and 2000 (au) in FSC were kept). After this FSC gating, we differentiated the accessory strain from the strain of interest using their mCerulean fluorescence. We considered events with BLU-V signal above 6·10^−2^ as cells from the accessory strain cells and events with BLU-V signal below 3·10^−2^ as cells from the strain of interest (Supplementary note 2). To convert raw cytometry data into fluorophore concentrations in relative promoter units^52^ (RPU), the fluorescence of each event in each channel was divided by its FSC to yield size-normalized fluorophore levels, and by the fluorescence of cells expressing the same fluorophore under the control of pTDH3. Finally, we subtract the mean fluorescence in RPU units during the two hours previous to light induction to all data points to avoid accounting for promoter leakage and autofluorescence. All the analysis for data processing is done using *Python 3*, with *pandas*^53^, *numpy*^54^, and *SciPy*^55^ packages.

### Secretion measurements and analysis

For measuring the secretion levels, we used 4% agarose micro-beads bound to the Anti-FLAG M2 monoclonal antibody (Anti-FLAG^®^ M2 Magnetic Beads from Sigma, M8823). Beads size ranges between 20-75 μm, which allowed us to use them in the cytometer. The protocol uses 0.5 μL of packed gel volume of beads per sample. After the beads equilibration (following the vendor recommended procedure), we collected the beads using the magnetic rack of the pipetting robot and resuspend them in 10 μL of water per assay. Then, 200 μL of culture samples were mixed with 26 μL of phosphate buffer 1 M and 10 μL of the equilibrated beads. In addition, all our beads measurements contained 26μL of mCerulean-3xFLAG at approximately 1 μg/ml (not used for analysis). After incubation during 1 hour at room temperature, the samples were washed three times before measurements by collecting the beads with the magnetic rack and resuspending them in 200 μL of TBS buffer. All manipulations with samples containing cells were done in the dark to avoid inducing the optogenetic system expressing the POI. Finally, bead solutions were passed to the cytometer and up to 1000 events were collected.

As for the yeast fluorescence measurements, we applied to beads the correction coefficients obtained using the *Guava*^®^ *easyCheck™* procedure. Despite the washing procedure, some cells were still present in the sample. We applied two gating conditions based on minimal side scatter (SSC), and minimal SSC/FSC scatter ratio. Then, to assess the secretion levels of the population in each reactor, we computed the median fluorescence on the GRN-B. To subtract autofluorescence and the residual crosstalk from the mCerulean-3xFLAG traces in the sample into the GRN-B channel, we used the beads corresponding to the non-induced sample as a blank. Thus, we subtract the median florescence of the non-induced sample to all the beads samples in each experiment. Additionally, since we used the accessory strain which abundance is changing in time, we normalized the secretion levels by the fraction of cells of interest in the population, corresponding to the fraction of cells that actually secrete. A label free proteomics analysis was used to obtain scaling factors for bead fluorescence to obtain measurements that are comparable for the different proteins (Supplementary note 4).

### Mathematical model

We used an ordinary differential equation model having three state variables and six parameters. Two parameters were fitted to data from cells expressing a non-secreted mNeonGreen protein and then kept constant. One parameter, a scaling factor for internal or secreted fluorescence values, was set based on secreted mNeonGreen data. The remaining three parameters were fitted on individual time series data for each induction level and each protein based on the non-accumulator cell population data. The non-accumulator and accumulator cell populations were distinguished using a Gaussian Mixture Model (Supplementary note 6, Python package *sklearn*.*mixture*.*GaussianMixture*^56^). The ODE model was integrated using the numerical solver *scipy*.*integrate*.*solve_ivp*^55^, and parameter fitting was performed thanks to the CMA-ES algorithm using the *pycma* package from Hansen and colleagues^57^.

## Acknowledgements

The authors would like to thank Chetan Aditya, Achille Fraisse, Allyson Holmes, Hélène Philippe, and Jakob Ruess for helpful discussions on cloning procedures or model development. We thank Mariette Matondo and Thibault Chaze of the proteomics facility of Institut Pasteur for LC-MS/MS protein quantification. This work was supported by ANR grants CyberCircuits (ANR-18-CE91-0002; S.S.-C.), MEMIP (ANR-16-CE33-0018, S.S.-C.), and SmartSec (ANR-21-CE44-0033, G.B.), by the H2020 Fet-Open COSY-BIO grant (grant agreement no. 766840, S.S.-C.) and by the Inria IPL grant COSY (G.B.).

## Author contributions

S.S.-C., F.B. and G.B. conceived the study. S.S.-C. constructed all strains, excepted the two *hac1* knock-out strains, performed all experiments, analyzed data, and developed mathematical models and control procedures. H.G. constructed the two *hac1* knock-out strains. S.N. and F.B. helped with software and hardware developments, and with tuning beads secretion assay protocols. F.B. and G.B. supervised the study. S.S.-C. and G.B. wrote the manuscript with input from all authors.

## Supplementary note 1. Heterologous proteins and yeast strains

Here we provide the list of proteins we studied and the strains we used in this work.

**Table S1.1.** Heterologous proteins of interest and some of their features

| Protein under study | POI name | PTM* | Native organism | Size (aa)** | GenBank ID |
| --- | --- | --- | --- | --- | --- |
| Non-secreted mNeonGreen<sup>1</sup> | Non-sec mNeon | N/A | N/A | 332 | BBB44438 |
| Secreted mNeonGreen<sup>1</sup> | mNeon | N/A | N/A | 354 | BBB44438 |
| Endo-1,4-beta-xylanase C<sup>2</sup> | XylC | 1x S-S | *Aspergillus niger* | 662 | EU848304 |
| Endo-1,4-beta-xylanase 2<sup>3</sup> | Xyl2 | 2x N-glyc | *Trichoderma reesei* | 545 | P36217.2 |
| Paraoxonase 1<sup>4</sup> | hPON1 | 1x S-S + 3x N-glyc | *Homo sapiens* | 709 | NM_000446.5 |
| Single chain variable fragment 4M5.3<sup>5</sup> | scFv | 2x S-S | N/A | 609 | 1X9Q_A |
| α-amylase<sup>6</sup> | Amy | 4x S-S + 2x N-glyc | *Aspergillus oryzae* | 832 | CAA31220 |

\* **PTM**: posttranslational modifications; **S-S**: disulfide bond; **N-glyc**: N-glycosylation. \*\* Includes, if present, the secretion tag, the protein of interest, the fluorescent reporter, and the three FLAG tag copies.

**Table S1.2.**
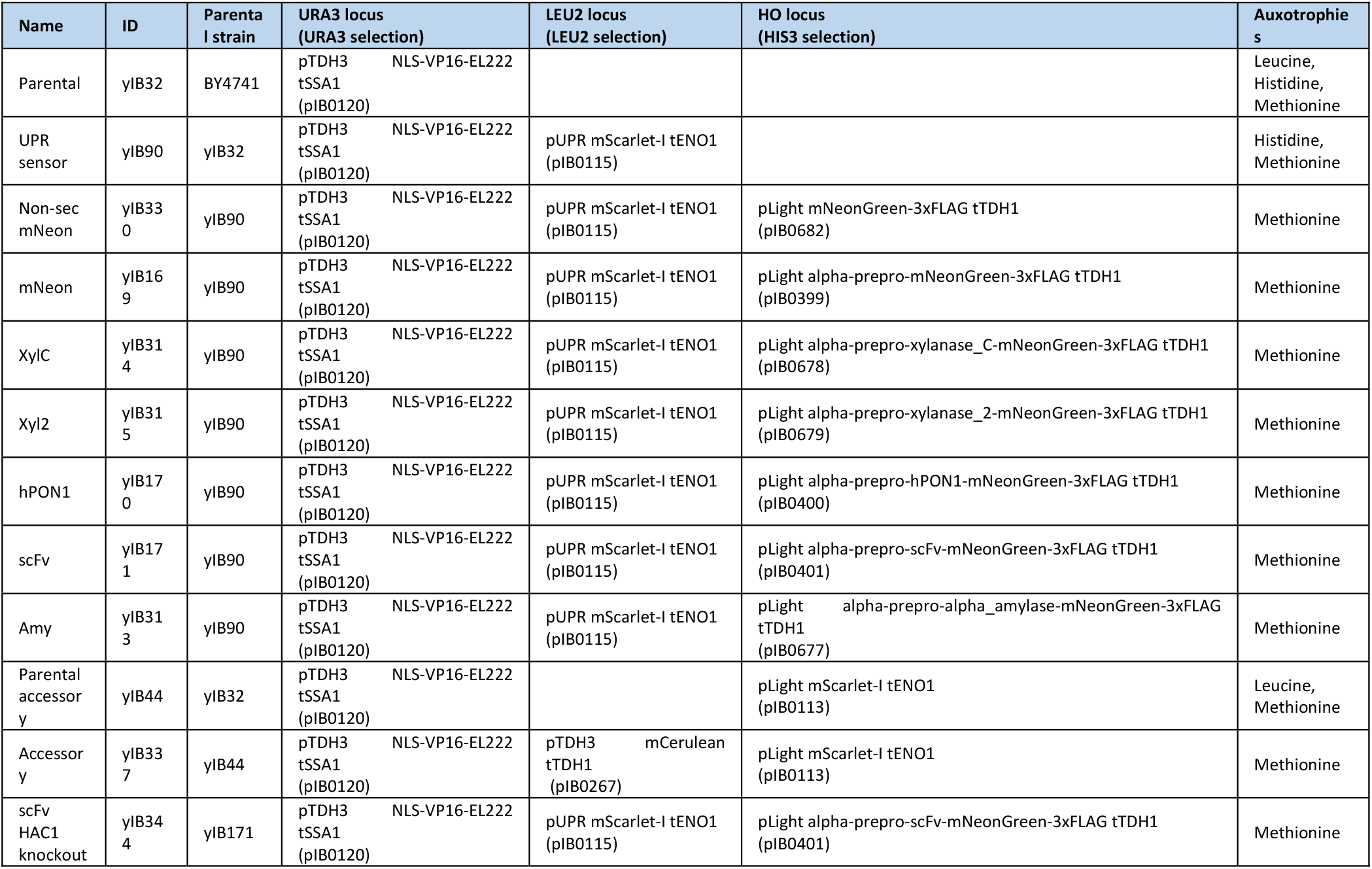
Description of the yeast strains used in this study

## Supplementary note 2. Use of an accessory strain

The accessory strain is co-cultured in all experiments (except the control experiments) at an initial ratio of 1:10 with respect to the strain of interest. It is used to guarantee a continuous flow of media through the reactors since its growth is not affected by the by secretory burden. Furthermore, it is also used to quantify actual induction demand in the strain of interest and to assess its growth rate.

### 2.1 Quantifying effective induction levels

The accessory strain has been constructed to achieve unbiased measurements of the induction levels actually perceived by cells. The accessory strain expresses a red fluorescent protein, mScarlet-I, whose expression is controlled by the EL222 optogenetic system as for the gene of interest in the strain under study. Therefore, at steady state, the expression levels of mScarlet-I in the accessory strain inform on the production rate of the POI in the strain under study.

**Fig. S2.1.**
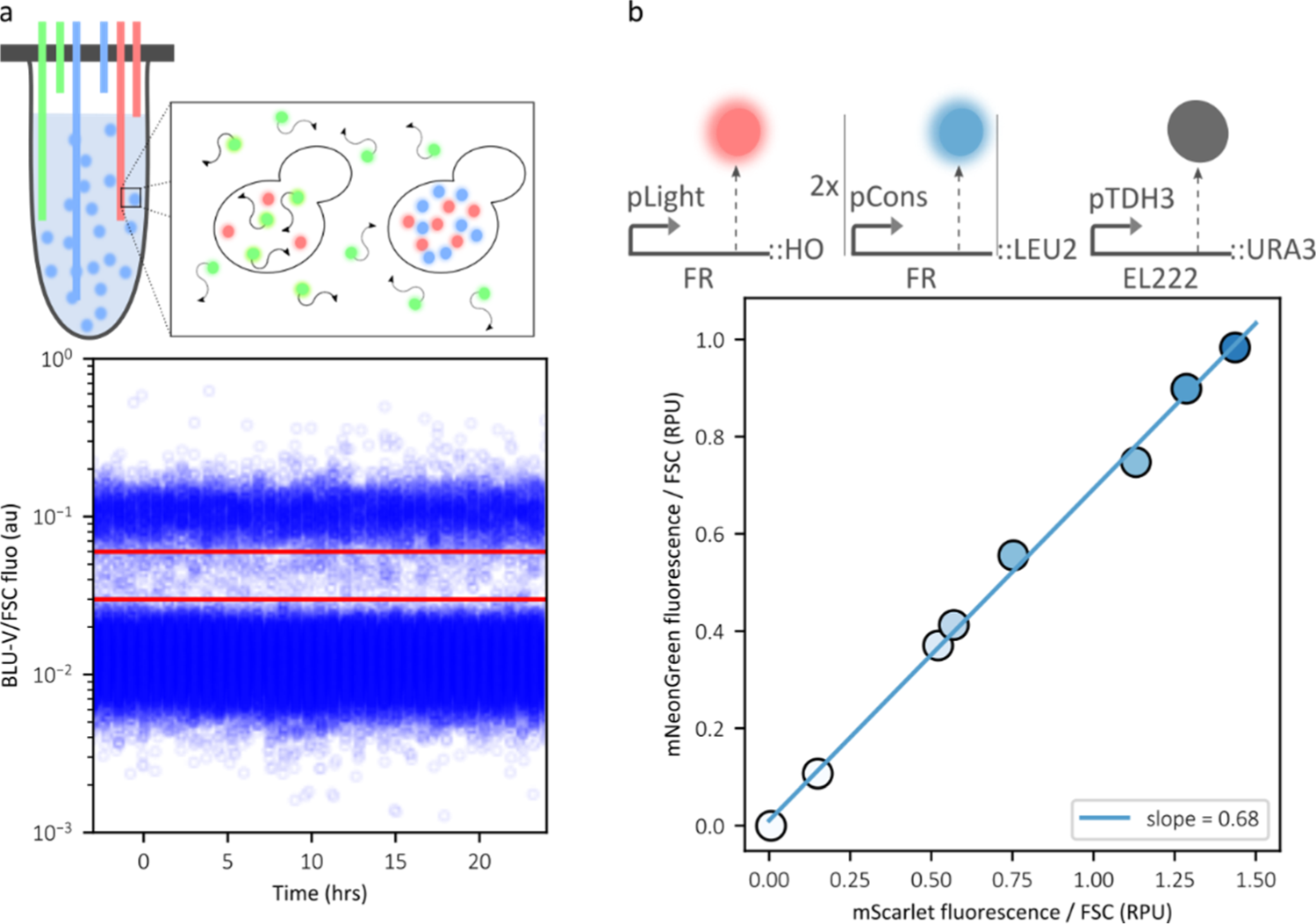
Use of the accessory strain to assess induction levels. **a** The accessory strain is co-cultured together with the strain of interest. It can be differentiated because it contains 2 copies of mCerulean under the control of a constitutive promoter (pTDH3 from *S. cerevisiae*). On the left, the plot shows an example of gating of the accessory strain co-cultured with the non-secreted mNeonGreen-expressing strain at maximal induction. The red solid lines represent the thresholds to differentiate each strain in the BLU-V channel (in this case normalized the FSC). Cells above the highest threshold are considered as accessory strain cells. Cells below the lowest threshold are considered as cells from the strain of interest. The events in between the two lines are not selected. **b** The accessory strain expresses a cytoplasmic red fluorescent protein, whose expression is controlled by the EL222 optogenetic system as for the gene of interest in the strain under study. The plot represents how the fluorescence detected for mScarlet-I in accessory strain and for mNeonGreen in the non-secreted mNeonGreen-expressing strain correlate. This confirms that the accessory cells can be used as a sensors of induction levels.

### 2.2 Assessing growth rate dynamics

When working in turbidostat mode, the OD is kept constant and one can in principle infer the growth rate from the influx rate of the supply pumps. These estimates were not very precise, however. Here, we use changes in the relative abundance of the accessory strain with respect to the strain of interest. By starting with a known fraction of accessory strain cells in the population, it is possible to assess growth rate of the strain under study by following the temporal evolution of the ratio of the strains, assuming that the growth rate of the accessory strain remains constant.

**Fig. S2.2.**
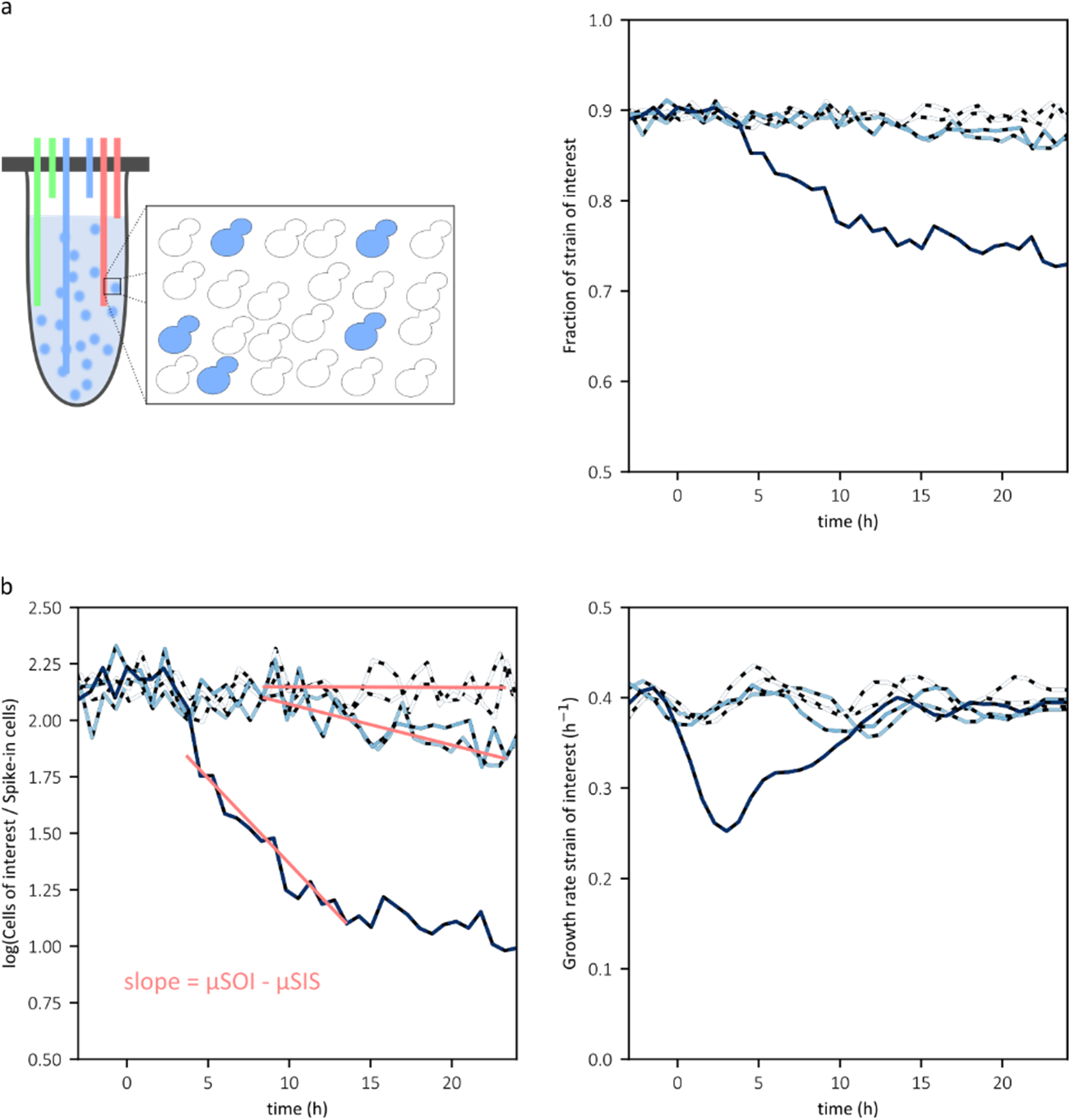
Use of the accessory strain to assess the growth rate of the strain of interest. **a** A scheme of the co-culture of accessory strain and strain under study in the bioreactor is shown on the left. The accessory strain produces mCerulean that is used to differentiate it from the strain of interest and monitor the dynamic changes of its fraction (right). **b** Computation of the growth rate of the strain of interest using the temporal evolution of the ratio of the two strains. μ: growth rate; SOI: strain of interest; SIS: accessory strain

## Supplementary note 3. iPOI location within the cell

With our constructs, the synthesized peptide is translocated to the ER after full translation. However, such process requires the protein to be unfolded, and therefore non-fluorescent. Consistently, we have observed by fluorescence microscopy that the non-secreted mNeonGreen is spread across the cytoplasm, whereas the secreted one is located exclusively in what should secretory compartments (ER, Golgi, vesicles, vacuole, etc).

**Fig. S3.1.**
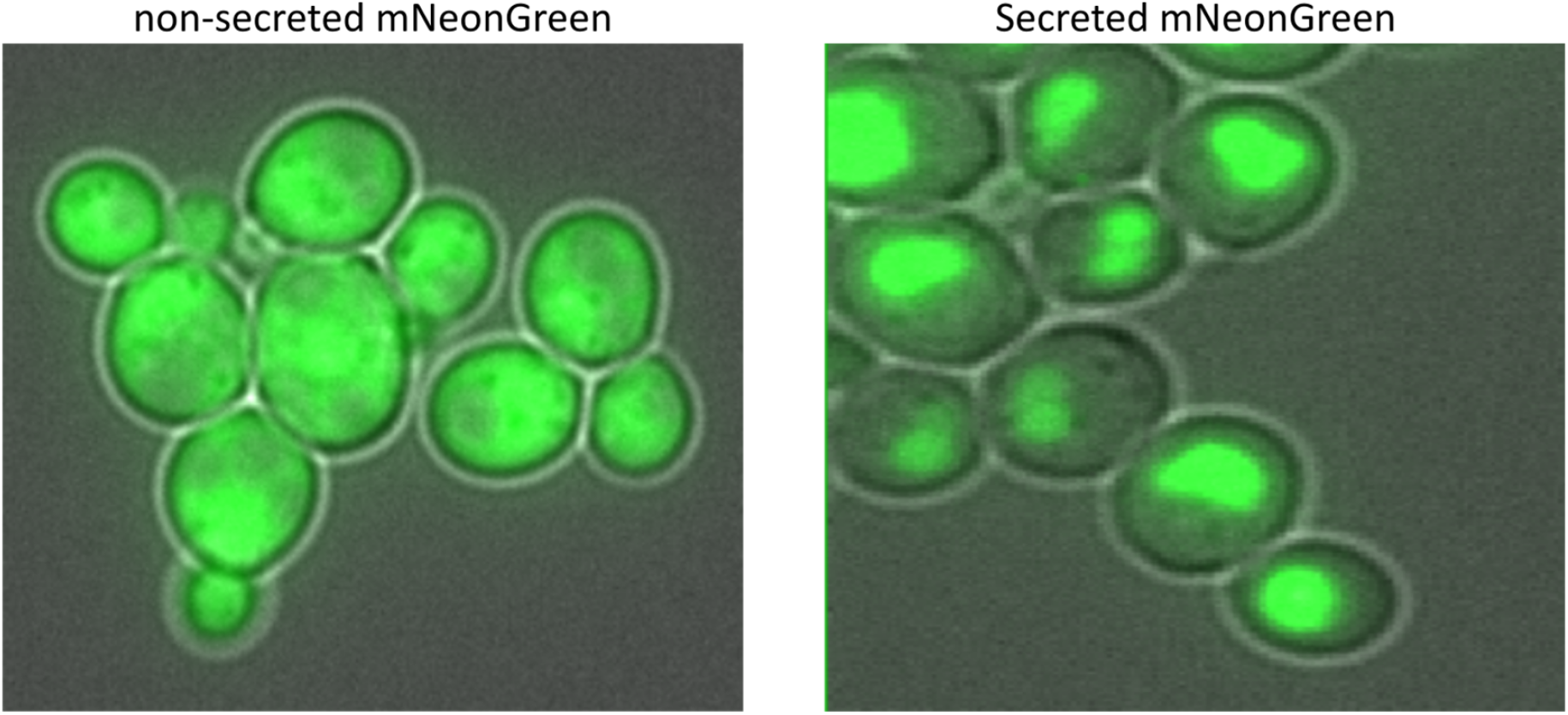
Fluorescence microscopy images of yeast cells producing mNeonGreen secreted and non secreted. On the left, cells expressing non-secreted mNeonGreen and on the right, cells expressing the secreted protein. In both cases the expression was induced by constant light.

## Supplementary note 4. Beads measurements

We provide details about the beads-based secretion measurements.

### 4.1 Secretion measurements principle

The goal is to develop a generic approach to detect secretion levels of a wide range of proteins using a cytometer. To gather secreted proteins together and so be detected by the cytometer, we developed a method in which agarose microbeads bound to antibodies recognize and bind a common epitope in all secreted POIs, the FLAG tag. By incubating a sample of the culture with the immuno-beads, only the secreted proteins have access to bind the beads, whereas those inside the cell cannot interact with the antibodies. The procedure works as follows: (i) a sample of the culture is incubated with the magnetic beads coated with the anti-FLAG antibody, (ii) using a magnetic grid and wash steps, the cells are separated from the beads and the beads can be passed through a flow cytometer.

### 4.2 Gating beads from residual cells

The distinction between cells and beads can be done by gating on the forward/side scatter pattern (FSC and SSC, respectively), since this property varies between cells and beads. The ratio SSC/FSC > 10 is used to distinguish beads from cells. We also gate to discard events having an SSC higher than the maximal SSC observed for cells (10^4^) (Figure S4.1).

For optimization of the method, different types and concentrations of beads were tested for sufficient binding properties and for not clogging the flow cytometer capillary. To test the binding of the beads and the feasibility of the method, the first assays were performed with pure commercial GFP fused to the FLAG tag at C-terminal in different concentrations (Figure S4.1). The results show a linear relationship between GFP concentrations and fluorescence readings on the cytometer, with the minimum concentration in the linear range being 0.02 nM, and the maximum being 90nM.

Then, we tested supernatants of batch culture after 24 hours of induction and containing residual cells, in which the concentrations of mNeonGreen were unknown. The fluorescence signal from the beads was proportional to the dilution factors of the supernatant in pure water. Finally, we checked whether this methodology is sensitive enough to measure the secretion levels from samples directly taken from our continuous culture bioreactors. To do so, we took a volume of the cell culture directly from the reactors at different levels of light induction and after 24 hours of induction. For further information in the beads measurements development check the thesis manuscript of Sebastian Sosa-Carrillo^7^.

**Fig. S4.1.**
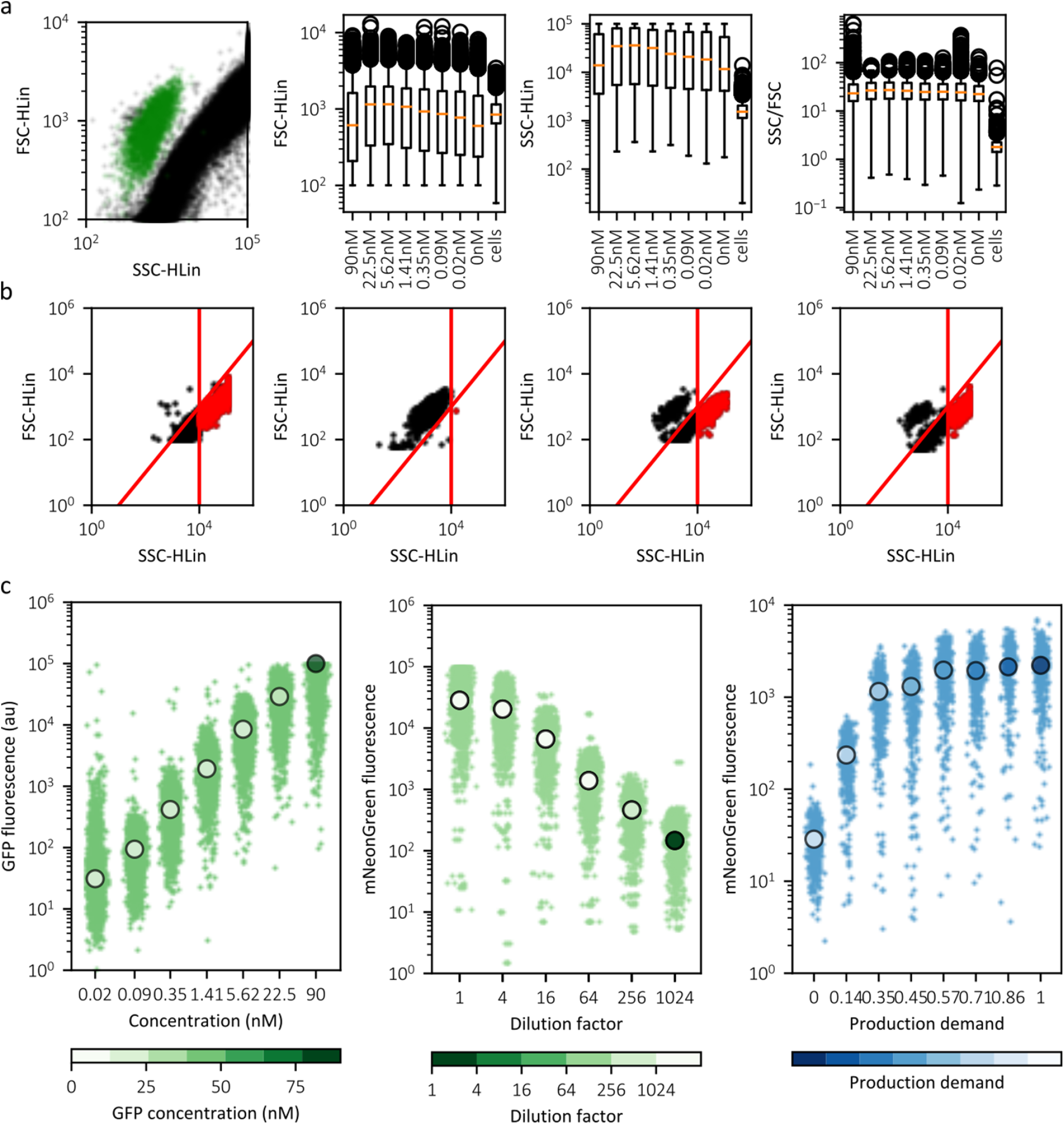
Characterization of the secretion measurements performed by using immuno-magnetic beads. **a** On left, we show the light scattering propertied of cells (green dots) and beads (black dots) from different experiments. On the right, the light scattering properties of beads with pure GFP and cells are represented in box plots. The data show that SSC and the ratio SSC/FSC are convenient for differentiation of both types of particles. **b** Examples of the gating strategy from samples of different nature. From left to right, a sample containing only beads, a sample containing only cells, a sample from a batch culture containing both beads and cells, a sample from a continuous culture from our setup containing both beads and cells. The two samples from cell cultures are shown after washing most of the cells by the protocol explained in materials and methods of the main text. The red lines indicate the gating criteria and the red dots are the gated events corresponding to beads. **c** Data obtained by the beads measurements protocol. On the left, the data obtained from the pure GFP assay, where GFP-FLAG has been incubated at different concentrations. The middle plot represented the data obtained from a batch culture after 24 hours of maximal induction to secrete mNeonGreen-3xFLAG. The right plot corresponds to data obtained from a continuous culture after 24 hours of maximal induction to secrete mNeonGreen-3xFLAG. In all cases the small dots correspond to the beads after gating, and the large dots to the median of the population.

### 4.3 Proteomics label-free quantification

We performed label-free quantitation of the POIs secreted from a continuous culture under full demand during 24 hours. Our goal was to define a coefficient to normalize the secretion levels measured with the beads for the possible differences in fluorescence produced by the structural context of the POI. We spiked 1 μl of the universal proteomics standard set (Sigma UPS2) into 7 μl of supernatants to assess the concentration of the POI. Thus, we obtained the iBAQ value, proportional to molar amount of peptides in the sample, of the full mixture and also coverage of the corresponding POIs in each sample. Then, knowing the different abundance of the UPS peptides in the mixture, we could assess the relative abundance of our POI (Figure S4.2, top). From that information we computed the relative difference on abundance of the different POIs in the samples, and we applied such coefficient to the obtained secretion levels measured by the beads methods (Figure S4.2, bottom and Table S4.1). The least abundant POIs (scFv) had a secretion level equal to one when it is expressed at full production demand during 24 hours. The rest of proteins are normalized in the same manner as shown in the following equation.

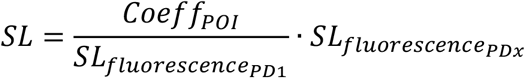

Each POI has a coefficient *Coeff*_*POI*_ that that is divided by the secretion levels obtained for such POI at production demand equal to one 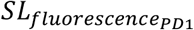, and then multiplied by the secretion level of all the production demand for that POI 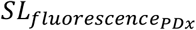.

**Table S4.1.** Coefficients used to normalize secretion levels

| Protein of interest | Name | Coeff | pmoles/ml |
| --- | --- | --- | --- |
| Secreted mNeonGreen<sup>1</sup> | mNeon | 11.6 | 47.5 |
| Endo-1,4-beta-xylanase C<sup>2</sup> | XylC | 12.6 | 51.7 |
| Endo-1,4-beta-xylanase 2<sup>3</sup> | Xyl2 | 3.4 | 14.1 |
| Paraoxonase 1<sup>4</sup> | hPON1 | 1.4 | 5.7 |
| Single chain variable fragment 4M5.3<sup>5</sup> | scFv | 1 | 4.1 |
| α-amylase<sup>6</sup> | amy | 2.3 | 9.3 |

**Fig. S4.2.**
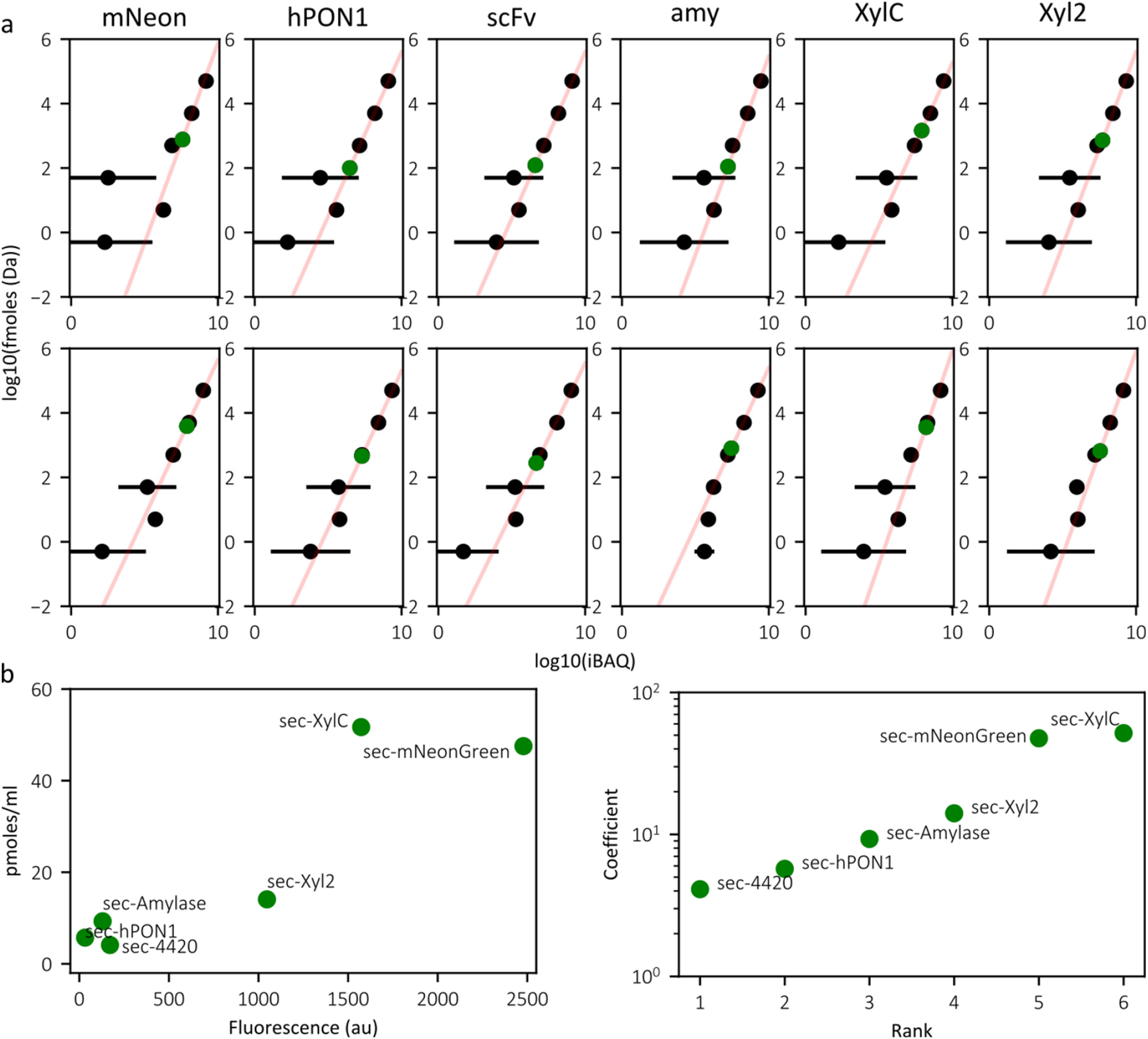
Label-free quantification using proteomics results. **a** Comparison of label-free quantification of UPS2 proteins (black points) of known molar quantity with the POI (green points) in different experiments. We performed linear regression (red line) using the UPS2 log10(iBAQ) of points between 6 and 10 to assess the molar quantity of the POI. **b** On the right, the plot shows the relationship between the inferred molar quantity (average of two replicas) and the fluorescence obtained from the beads. On the left, we represent the coefficients for each of the POI that are used to normalize the fluorescence values from beads and obtain the secretion levels shown in this study.

## Supplementary note 5. Growth rate analysis

We provide details on the growth rates characterization performed in this study and its relation with the presence of accumulators cells.

### 5.1 Reference growth rate

The growth rate used as reference to compute the growth rate of the strain of interest as explained in Supplementary note 2 is the one obtained from the accessory strain. It was computed from the optical density at 600nm (OD_600_), controlled to remain between 0.4 and 0.6 by the bioreactors platform.

**Fig. S5.1.**
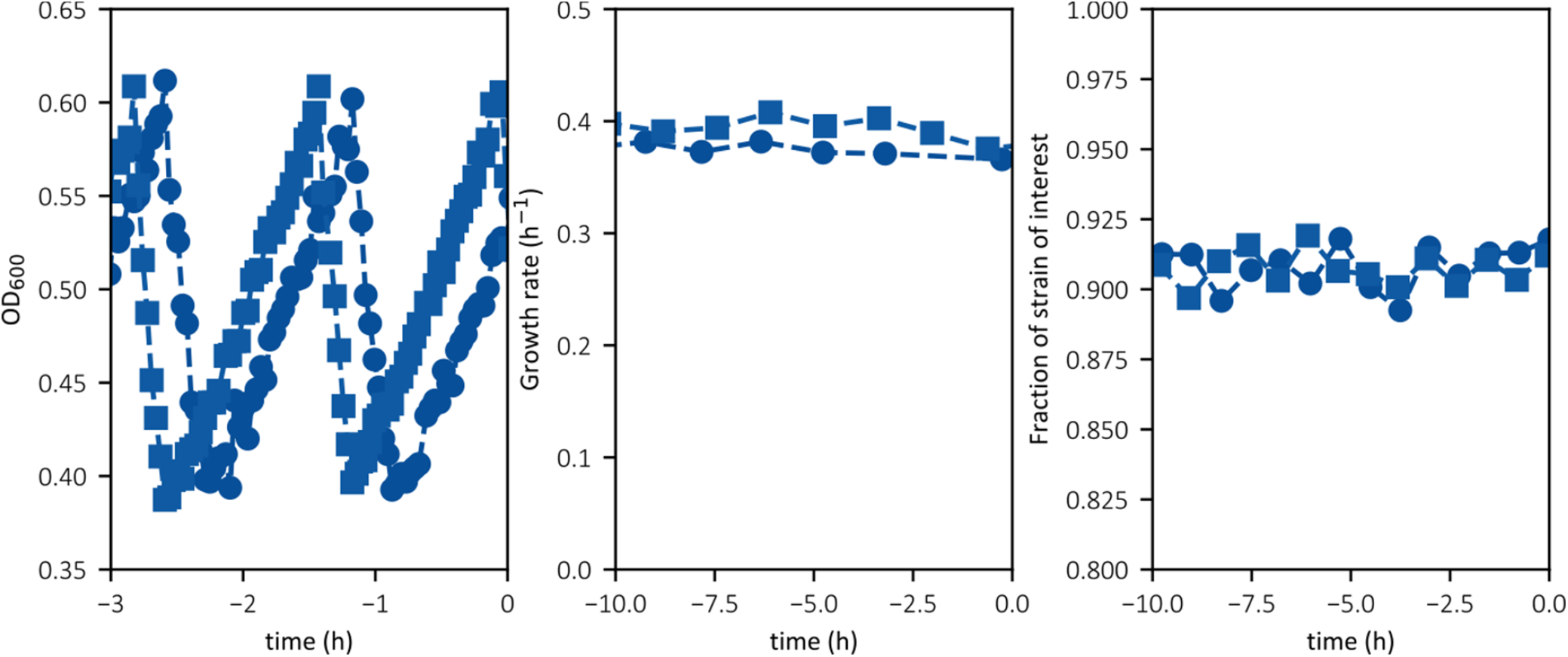
Absolute growth rate assessment from a co-culture. The plot on the left shows the OD oscillating between the two ODs set at 0.4 and 0.6. The data is obtained from two replicas of a co-culture of accessory strain and scFv-secreting cells, non-induced, growing in standard conditions. The plot in the middle shows the computed growth rate from the OD increase by doing linear regression on the logarithm. The plot on the right shows the proportion of each of the strain in the co-culture prior to induction to show that no growth decay is observed in strains.

### 5.2 Growth rates in characterization experiments

Here we show the growth rate for the characterization experiments shown in the main text for each of the proteins.

**Fig. S5.2.**
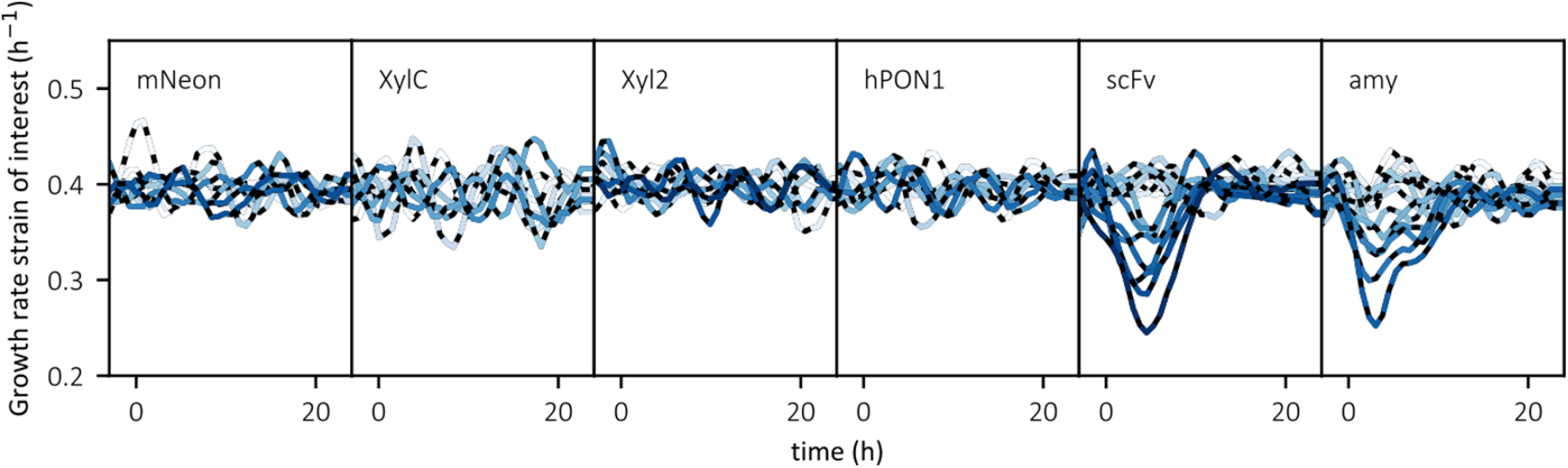
Computed growth rate in characterization experiments. Each plot shows the growth rate corresponding to each strain of interest during the characterization experiments (from 3 hours prior to induction to 24 hours after induction) for all the different production demands. The intensity of the blue color corresponds to the induction levels.

### 5.3 The fraction of accumulator cells is correlated in time with the overall growth rate

The growth rate of the population decays proportionally to the fraction of accumulators, indicating that accumulator cells have a strongly reduced growth rate. Moreover, eventually all accumulator cells defined by their iPOI levels also become accumulators as defined by their UPR levels.

**Fig. S5.3.**
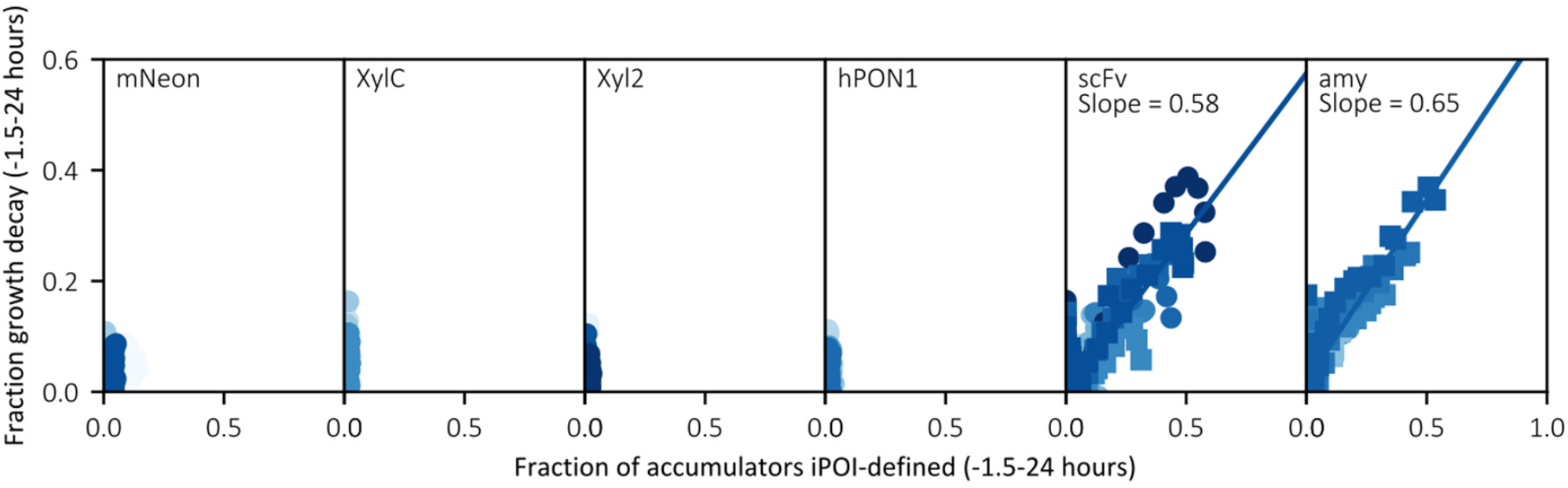
Relation between the fraction of accumulators and the population growth decay. Each plot shows the fraction of growth rate decay as a function of the fraction of the accumulator cells corresponding to each strain of interest during the characterization experiments (from 1.5 hours prior to induction to 24 hours after induction) for all the different production demands. The intensity of the blue color corresponds to the strength of the production demand. The slopes indicated for scFv and α-amylase-secreting cells have been computed by linear regression. In this case the fraction of accumulators is computed from the overall population, including the accessory strain.

## Supplementary note 6. Accumulator population analysis

We provide details on the characterization of the accumulator cells performed in this study.

### 6.1 iPOI levels in accumulator cells increases proportionally to induction levels

The iPOI levels for accumulator cells increase linearly with the induction levels. This is not the case for non-accumulator cells. This observation, together with the plateau observed in the secretion levels, indicates that the pathway capacity of the cell is saturated for both degradation and secretion.

**Fig. S6.1.**
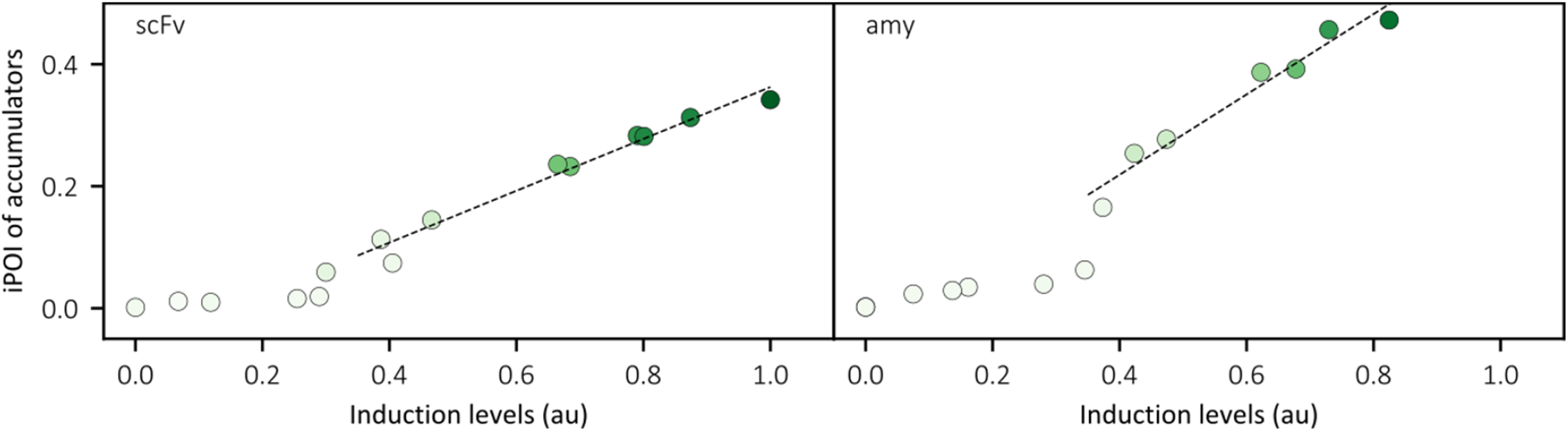
Relation between iPOI and induction levels in accumulator cells. Each plot shows the increase in iPOI levels with induction levels for the accumulator cells. The intensity of the green color is proportional to the fraction of accumulators. The dashed black line corresponds to a linear fit starting with induction levels above 0.35 to 1.

### 6.2 Differentiating between accumulator and non-accumulator cells

To distinguish the two populations, we used a Gaussian mixture model provided in the Python package sklearn.mixture.GaussianMixture.

**Fig. S6.2.**
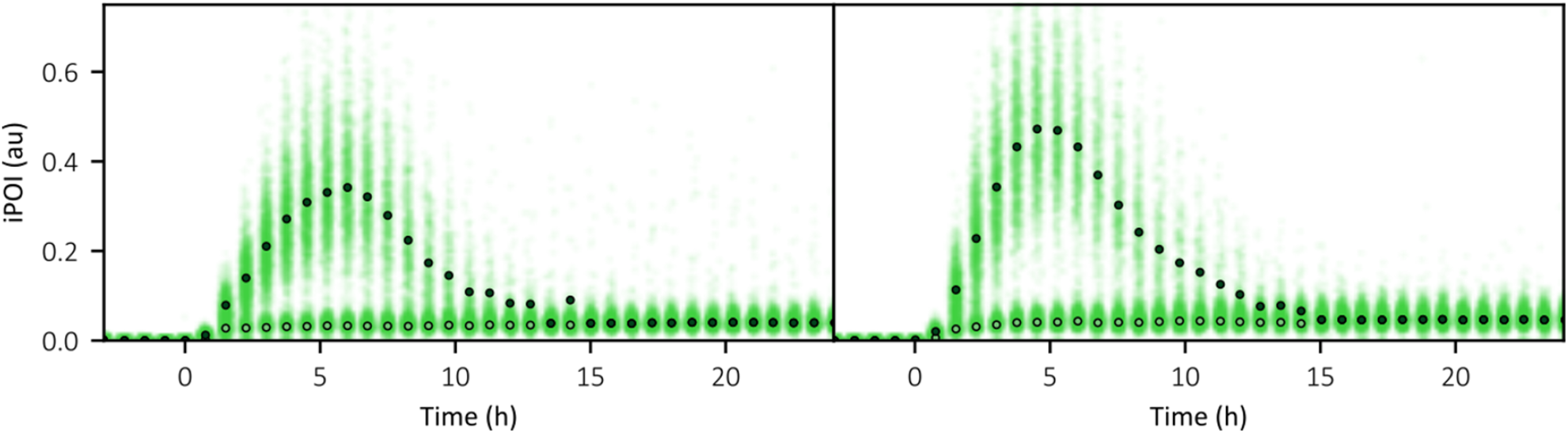
Defining the accumulator and non-accumulator populations. The plot shows the two populations in two experiments. In each of them the temporal evolution of iPOI is shown over time. The median for accumulators at each time point is represented as a black dot, the median for non-accumulators as a white dote. The left plot corresponds to the experiment of scFv-secreting cells at maximal induction levels. The right plot corresponds to the experiment of amylase-secreting cells at maximal induction levels.

### 6.3 Correlation of the maximal fractions of iPOI- and UPR-defined accumulators

The maximal fraction of accumulators defined by their iPOI levels is correlated with the maximal fraction of accumulators defined by their UPR levels. The difference between the two maxima is the time at which they are reached.

**Fig. S6.3.**
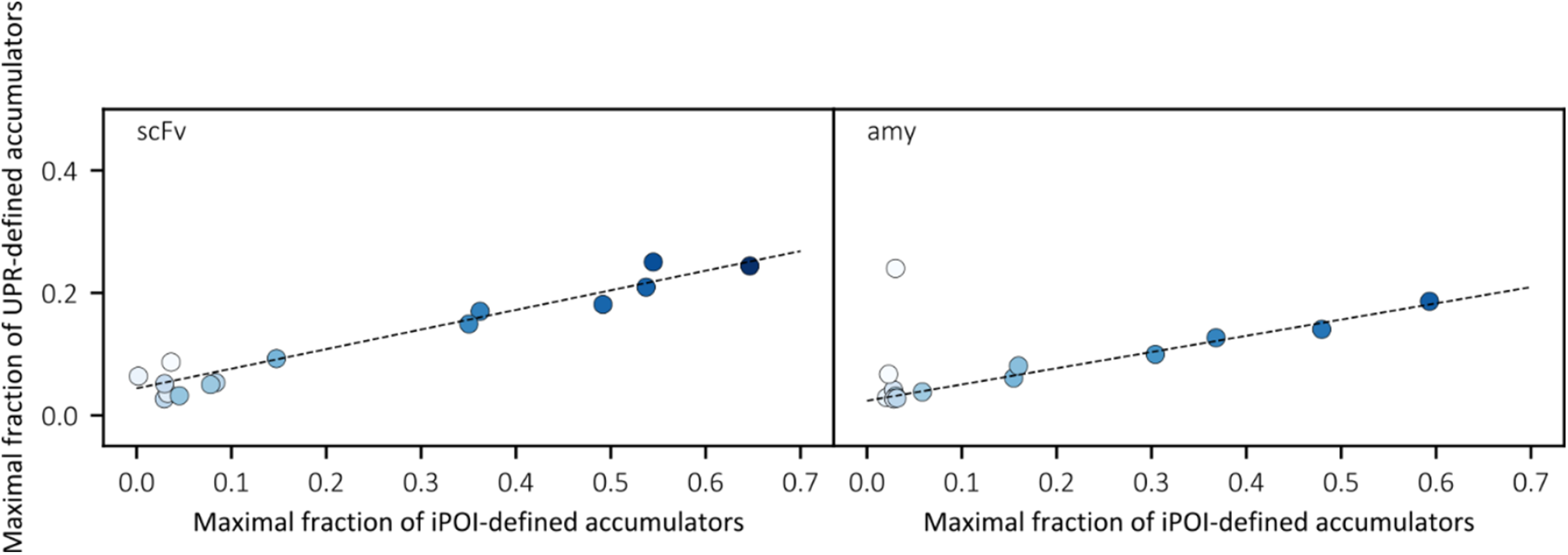
Correlation between the maximal fractions of accumulators defined by their iPOI or UPR levels. In each plot the intensity of the blue color is proportional to the induction levels. The black line indicates the linear relation between the two variables.

## References

1. Yu, L. P., Wu, F. Q. & Chen, G. Q. Next-Generation Industrial Biotechnology-Transforming the Current Industrial Biotechnology into Competitive Processes. Biotechnology Journal 14, 1–9 (2019).

2. Campbell, K., Xia, J. & Nielsen, J. The Impact of Systems Biology on Bioprocessing. Trends in Biotechnology 35, 1156–1168 (2017).

3. Wang, G., Huang, M. & Nielsen, J. Exploring the potential of Saccharomyces cerevisiae for biopharmaceutical protein production. Current Opinion in Biotechnology 48, 77–84 (2017).

4. Walsh, G. Biopharmaceutical benchmarks 2018. Nature Biotechnology 36, 1136–1145 (2018).

5. Mattanovich, D., Sauer, M. & Gasser, B. Yeast biotechnology: teaching the old dog new tricks. (2014) doi:10.1186/1475-2859-13-34.

6. Buchberger, A., Bukau, B. & Sommer, T. Protein Quality Control in the Cytosol and the Endoplasmic Reticulum: Brothers in Arms. Molecular Cell 40, 238–252 (2010).

7. Delic, M. et al. The secretory pathway: Exploring yeast diversity. FEMS Microbiology Reviews 37, 872–914 (2013).

8. Lee, S. Y. & Kim, H. U. Systems strategies for developing industrial microbial strains. Nature Biotechnology 33, 1061–1072 (2015).

9. Tang, H. et al. Engineering protein folding and translocation improves heterologous protein secretion in Saccharomyces cerevisiae. Biotechnology and Bioengineering 112, 1872–1882 (2015).

10. Thak, E. J., Yoo, S. J., Moon, H. Y. & Kang, H. A. Yeast synthetic biology for designed cell factories producing secretory recombinant proteins. FEMS Yeast Research 20, 1–17 (2020).

11. Travers, K. J. et al. Functional and genomic analyses reveal an essential coordination between the unfolded protein response and ER-associated degradation. Cell 101, 249–258 (2000).

12. Hampton, R. Y. ER stress response: Getting the UPR hand on misfolded proteins. Current Biology 10, 518–521 (2000).

13. Halbleib, K. et al. Activation of the Unfolded Protein Response by Lipid Bilayer Stress. Molecular Cell 67, 673-684.e8 (2017).

14. Peter, W. & David, R. The Unfolded Protein Response. Science 1, 1–3 (2011).

15. Ng, D. T. W., Spear, E. D. & Walter, P. The unfolded protein response regulates multiple aspects of secretory and membrane protein biogenesis and endoplasmic reticulum quality control. Journal of Cell Biology 150, 77–88 (2000).

16. Brodsky, J. L. & McCracken, A. A. ER protein quality control and proteasome-mediated protein degradation. Seminars in Cell & Developmental Biology 10, 507–513 (1999).

17. Bernales, S., McDonald, K. L. & Walter, P. Autophagy counterbalances endoplasmic reticulum expansion during the unfolded protein response. PLoS Biology 4, 2311–2324 (2006).

18. Song, S., Tan, J., Miao, Y. & Zhang, Q. Crosstalk of ER stress-mediated autophagy and ER-phagy: Involvement of UPR and the core autophagy machinery. Journal of Cellular Physiology 233, 3867–3874 (2018).

19. Lajoie, P. & Snapp, E. L. Size-dependent secretory protein reflux into the cytosol in association with acute endoplasmic reticulum stress. Traffic 21, 419–429 (2020).

20. Igbaria, A. et al. Chaperone-mediated reflux of secretory proteins to the cytosol during endoplasmic reticulum stress. Proceedings of the National Academy of Sciences 116, 11291–11298 (2019).

21. Bertaux, F. et al. Enhancing bioreactor arrays for automated measurements and reactive control with ReacSight. Nature Communications 2022 13:1 13, 1–12 (2022).

22. Brake, A. J. et al. α-Factor-directed synthesis and secretion of mature foreign proteins in Saccharomyces cerevisiae. Proceedings of the National Academy of Sciences of the United States of America 81, 4642–4646 (1984).

23. Liu, Z., Tyo, K. E. J., Martínez, J. L., Petranovic, D. & Nielsen, J. Different expression systems for production of recombinant proteins in Saccharomyces cerevisiae. Biotechnology and Bioengineering 109, 1259–1268 (2012).

24. Shaner, N. C. et al. A bright monomeric green fluorescent protein derived from Branchiostoma lanceolatum. Nature Methods 2013 10:5 10, 407–409 (2013).

25. Motta-Mena, L. B. et al. An optogenetic gene expression system with rapid activation and deactivation kinetics. Nature Chemical Biology 10, 196–202 (2014).

26. Pincus, D. et al. BiP binding to the ER-stress sensor Ire1 tunes the homeostatic behavior of the unfolded protein response. PLoS Biology 8, (2010).

27. Bindels, D. S. et al. mScarlet: a bright monomeric red fluorescent protein for cellular imaging. Nature Methods 2016 14:1 14, 53–56 (2016).

28. Davidson, E. A., Basu, A. S. & Bayer, T. S. Programming Microbes Using Pulse Width Modulation of Optical Signals. Journal of Molecular Biology 425, 4161–4166 (2013).

29. Benzinger, D. & Khammash, M. Pulsatile inputs achieve tunable attenuation of gene expression variability and graded multi-gene regulation. Nature Communications 1–38 (2018) doi:10.1038/s41467-018-05882-2.

30. Laboissiere, M. C. A., Sturley, S. L. & Raines, R. T. The Essential Function of Protein-disulfide Isomerase Is to Unscramble Non-native Disulfide Bonds (*). Journal of Biological Chemistry 270, 28006–28009 (1995).

31. Fu, J., Gao, J., Liang, Z. & Yang, D. PDI-Regulated Disulfide Bond Formation in Protein Folding and Biomolecular Assembly. Molecules 2021, Vol. 26, Page 171 26, 171 (2020).

32. Babour, A., Bicknell, A. A., Tourtellotte, J. & Niwa, M. A Surveillance Pathway Monitors the Fitness of the Endoplasmic Reticulum to Control Its Inheritance. Cell 142, 256–269 (2010).

33. Jonas, F. R. H., Royle, K. E., Aw, R., Stan, G. B. V. & Polizzi, K. M. Investigating the consequences of asymmetric endoplasmic reticulum inheritance in Saccharomyces cerevisiae under stress using a combination of single cell measurements and mathematical modelling. Synthetic and Systems Biotechnology 3, 64–75 (2018).

34. Kafri, M., Metzl-Raz, E., Jona, G. & Barkai, N. The Cost of Protein Production. Cell Reports 14, 22–31 (2016).

35. Metzl-Raz, E. et al. Principles of cellular resource allocation revealed by condition-dependent proteome profiling. eLife 6, 1–21 (2017).

36. Cox, J. S. & Walter, P. A Novel Mechanism for Regulating Activity of a Transcription Factor That Controls the Unfolded Protein Response. Cell 87, 391–404 (1996).

37. Wang, Y. et al. Precision and functional specificity in mRNA decay. PNAS April 30, 5860–5865 (2002).

38. Geisberg, J. V., Moqtaderi, Z., Fan, X., Ozsolak, F. & Struhl, K. Global Analysis of mRNA Isoform Half-Lives Reveals Stabilizing and Destabilizing Elements in Yeast. Cell 156, 812–824 (2014).

39. Aditya, C., Bertaux, F., Batt, G. & Ruess, J. A light tunable differentiation system for the creation and control of consortia in yeast. Nature Communications 2021 12:1 12, 1–10 (2021).

40. Aditya, C., Bertaux, F., Batt, G. & Ruess, J. Using single-cell models to predict the functionality of synthetic circuits at the population scale. Proceedings of the National Academy of Sciences of the United States of America 119, e2114438119 (2022).

41. Love, K. R. et al. Systematic single-cell analysis of pichia pastoris reveals secretory capacity limits productivity. PLoS ONE 7, 1–11 (2012).

42. Wittrup, K. D., Robinson, A. S., Parekh, R. N. & Forrester, K. J. Existence of an optimum expression level for secretion of foreign proteins in yeast. Annals of the New York Academy of Sciences 745, 321–330 (1994).

43. Parekh, R. N. & Wittrup, K. D. Expression level tuning for optimal heterologous protein secretion in Saccharomyces cerevisiae. Biotechnology Progress 13, 117–122 (1997).

44. Lee, M. E., DeLoache, W. C., Cervantes, B. & Dueber, J. E. A Highly Characterized Yeast Toolkit for Modular, Multipart Assembly. ACS Synthetic Biology 4, 975–986 (2015).

45. Benson, D. A. et al. GenBank. Nucleic Acids Research 43, D30 (2015).

46. Gietz, R. D. Yeast transformation by the LiAc/SS carrier DNA/PEG method. Methods in molecular biology (Clifton, N.J.) 1205, 1–12 (2014).

47. Gietz, R. D. & Woods, R. A. Transformation of yeast by lithium acetate/single-stranded carrier DNA/polyethylene glycol method. Methods in Enzymology 350, 87–96 (2002).

48. Agier, N., Fleiss, A., Delmas, S. & Fischer, G. A versatile protocol to generate translocations in yeast genomes using crispr/cas9. Methods in Molecular Biology 2196, 181–198 (2021).

49. Chung, B. H. & Park, K. S. Simple Approach to Reducing Proteolysis During Secretary Production of Human Parathyroid Hormone in Saccharomyces cerevisiae. (1998).

50. Kang, H. A. et al. Proteolytic stability of recombinant human serum albumin secreted in the yeast Saccharomyces cerevisiae. Applied Microbiology and Biotechnology 53, 575–582 (2000).

51. Peña, A., Sánchez, N. S., Álvarez, H., Calahorra, M. & Ramírez, J. Effects of high medium pH on growth, metabolism and transport in Saccharomyces cerevisiae. FEMS Yeast Research 15, 5 (2015).

52. Kelly, J. R. et al. Measuring the activity of BioBrick promoters using an in vivo reference standard. Journal of Biological Engineering 3, 1–13 (2009).

53. McKinney, W. Data Structures for Statistical Computing in Python. in Proceedings of the 9th Python in Science Conference 56–61 (2010). doi:10.25080/majora-92bf1922-00a.

54. Harris, C. R. et al. Array programming with NumPy. Nature 585, 357 (2020).

55. Virtanen, P. et al. SciPy 1.0: fundamental algorithms for scientific computing in Python. Nature Methods 2020 17:3 17, 261–272 (2020).

56. Pedregosa, F. et al. Scikit-learn: Machine Learning in Python. Journal of Machine Learning Research 12, 2825–2830 (2011).

57. Hansen, N. A global surrogate assisted CMA-ES. GECCO 2019 -Proceedings of the 2019 Genetic and Evolutionary Computation Conference 664–672 (2019) doi:10.1145/3321707.3321842.

## References

1. Shaner, N. C. et al. A bright monomeric green fluorescent protein derived from Branchiostoma lanceolatum. Nature Methods 2013 10:5 10, 407–409 (2013).

2. Do, T. T., Quyen, D. T., Nguyen, T. N. & Nguyen, V. T. Molecular characterization of a glycosyl hydrolase family 10 xylanase from Aspergillus niger. Protein Expression and Purification 92, 196–202 (2013).

3. Törrönen, A. et al. The Two Major Xylanases from Trichoderma Reesei: Characterization of Both Enzymes and Genes. Bio/Technology 1992 10:11 10, 1461–1465 (1992).

4. Ünver, Y., Kurbanoğlu, E. B. & Erdoğan, O. Expression, purification, and characterization of recombinant human paraoxonase 1 (rhPON1) in Pichia pastoris. Turkish Journal of Biology 39, 649–655 (2015).

5. Boder, E. T., Midelfort, K. S. & Wittrup, K. D. Directed evolution of antibody fragments with monovalent femtomolar antigen-binding affinity. Proceedings of the National Academy of Sciences of the United States of America 97, 10701–10705 (2000).

6. Liu, Z., Tyo, K. E. J., Martínez, J. L., Petranovic, D. & Nielsen, J. Different expression systems for production of recombinant proteins in Saccharomyces cerevisiae. Biotechnology and Bioengineering 109, 1259–1268 (2012).

7. Sosa-Carrillo, S. Pipeline for the systematic characterization of heterologous protein secretory load to assess bioproduction efficiency. https://hal.inria.fr/tel-03612661/.

